# The specific interface for selenocysteine-decoding tRNA in HIV-1 Gag MA domain regulates progeny virion infectivity

**DOI:** 10.64898/2026.10.16.759769

**Authors:** Rajat Mudgal, Dishari Thornhill, Kai-Neng Chou, Hamza Qureshi, Akira Ono

## Abstract

Localization of HIV-1 particle assembly at the plasma membrane (PM) is mediated by the HIV-1 Gag matrix (MA) domain. Competing binding of PM acidic phospholipids and cellular tRNAs to MA-highly basic region (MA-HBR) likely regulate the PM-specific binding of Gag; however, the specificity of the MA-tRNA interaction and its role in viral replication remain poorly understood. Here, using a modified crosslinking-immunoprecipitation (CLIP)-sequencing workflow for identifying MA-bound tRNAs, we determined that most MA-bound tRNAs were also abundantly present in cells. However, selenocysteine-tRNA (SeCTCA) is overrepresented among Gag-bound tRNAs despite its paucity. The binding of SeCTCA to Gag is entirely dependent on MA-HBR, whereas non-SeCTCA tRNAs can bind Gag through both MA and nucleocapsid domains. Structure prediction of MA-tRNA complexes revealed the binding interface unique to the MA-SeCTCA complex, which was verified experimentally. CLIP experiments demonstrated that this interface is specifically needed for the MA binding to SeCTCA but not to other tRNAs. Notably, disruption of this interface altered Gag localization pattern, reduced infectivity of virions, and impaired virus replication in CD4+ T cells. Collectively, our results indicate that SeCTCA tRNA binds the HIV-1 Gag MA domain with high specificity and promotes virion formation and efficient replication in T cells.

**Teaser:** Among tRNAs that bind HIV structural protein Gag, selenocysteine tRNA does so in a unique way, which facilitates infection

## INTRODUCTION

Assembly of the HIV-1 takes place at the plasma membrane (PM) for most cell types including CD4^+^ T cells, the physiological host cell type for HIV-1 (*1–5*). The assembly process is driven by the viral structural polyprotein Gag. Gag has four major domains: the matrix (MA) domain mediates targeting and anchoring of Gag to the PM, the capsid (CA) domain facilitates Gag multimerization via protein-protein interactions, the nucleocapsid (NC) domain selectively packages viral RNA while also contributing to Gag multimerization, and p6 recruits the cellular endosomal sorting complex required for transport for the release of the budding viral particle from the PM (*1*).

MA and NC are the two major RNA binding domains of Gag. They bind RNA with different specificities. NC mediates packaging of viral genomic RNA but otherwise primarily binds host mRNAs (*6–10*). MA has been shown to bind various RNAs, in particular tRNAs, in previous in vitro and cell-based studies (*9*, *11–15*). The globular head of MA is myristoylated, which drives the stable binding of Gag to the PM (*16*, *17*). The myristate moiety is sequestered in a hydrophobic pocket of the globular head in the cytosol, but upon multimerization at the lipid bilayer, it is exposed and is inserted into the membrane (*18*). The other signal in the MA globular head that mediates membrane binding is the highly basic region (HBR), which forms the binding site for negatively charged phospholipids (*19–23*). RNase treatment of Gag produced in rabbit reticulocyte lysates drastically promotes its binding to liposomes consisting of phosphatidylcholine (PC) and phosphatidylserine (PS) (PC+PS), whereas the effect of RNase is limited when liposomes contain phosphatidylinositol-4,5-bisphosphate (PIP2), a phospholipid found in the inner leaflet of PM (*13*, *24*, *25*). Therefore, the binding of RNA to Gag has been proposed to suppress the electrostatic binding to membranes that do not contain PIP2, thereby ensuring preferential localization of Gag to the PM. Notably, MA with HBR mutations that severely disrupts MA-RNA binding showed promiscuous localization of Gag to different cellular membrane compartments (*26*). Although NC also binds to RNA in the cells, inhibition of Gag membrane binding by RNA is attributed to MA-RNA interaction, as RNase treatment of the Gag variant lacking the NC domain (delNC) still led to an increase in binding to PC+PS liposome membranes (*13*, *24*, *27*).

A photoactivatable ribonucleoside-enhanced crosslinking and immunoprecipitation RNA sequencing (PARCLIP-RNA-seq) study using HIV-1-expressing HEK293T cells demonstrated that RNAs bound to MA in cells are primarily tRNAs, especially a subset of tRNAs decoding Gly, Glu, Lys, and Val (*9*). In in vitro assays, addition of yeast tRNAs inhibits binding of HIV-1 Gag to PC+PS liposomes but not to liposomes containing PIP2, indicating that PIP2 can overcome the Gag-tRNA interaction (*27*, *28*). Previous studies have also shown that a Lys tRNA (LysTTT) inhibits MA membrane binding more strongly than Pro tRNA in in vitro liposome binding assays (*28*). Further, RNA sequencing, structural, and mutagenesis analyses indicated that the elbow region, especially the D arm and T arm, of LysTTT is essential for binding to Gag or inhibition of Gag-membrane binding (*9*, *28*, *29*). Differential inhibition of Gag-membrane binding by different tRNAs and the dependence on the specific tRNA regions for MA binding of tRNA suggest that sequence/structural features of tRNA are critical determinants for inhibition of Gag binding to non-PIP2-containing membranes and hence for the PM-specific and/or timely virus assembly (*9*, *13*, *24*, *27–29*). However, molecular mechanisms determining the binding specificity of Gag for tRNAs as well as the effects of Gag-membrane binding on Gag-tRNA binding remain to be fully elucidated in cell-based analyses.

The current model for tRNA-mediated regulation of Gag membrane targeting suggests that binding of Gag to the PM leads to the dissociation of Gag-bound tRNA, since binding sites for PIP2 and tRNA partially overlap with each other at MA-HBR (*13*, *22*, *23*, *26*, *29–31*). The previous PARCLIP-RNA-seq study examined the RNAs cross-linked to Gag that is competent in membrane binding. However, according to the model above, membrane binding and tRNA binding of MA are mutually exclusive. Therefore, tRNAs recovered with membrane binding-competent Gag would not represent the entire population of tRNAs that can bind MA. Additionally, it remains to be determined whether Gag preference for some tRNAs observed in the previous PARCLIP study might simply reflect the cytosolic abundance of the individual tRNAs as more abundant tRNAs would have a greater chance to encounter a Gag molecule and bind it.

To define the tRNA binding specificity of MA-HBR, using a modified PARCLIP-RNA-seq approach, we compared the abundances of tRNAs cross-linked to a delNC Gag construct that does not have a myristoylation site (non-myr) and hence is membrane binding-incompetent (non-myr delNC Gag) in the cytoplasm versus total tRNAs present in the cells. Most tRNAs bound to delNC Gag, including LysTTT, were tRNAs abundantly present in cells. However, despite its low cellular levels, selenocysteine-decoding tRNA (SeCTCA) was found in the non-myr delNC Gag-bound pool of tRNAs with high frequency. The PARCLIP-RT-qPCR analysis indicated that most tRNAs did not require intact MA-HBR for binding to full-length Gag containing the NC domain, whereas SeCTCA was strictly dependent on MA-HBR. Furthermore, the in vitro liposome binding assay showed that SeCTCA inhibits Gag membrane binding with significantly higher efficiency than LysTTT. Finally, we found that MA-HBR forms a unique interface with SeCTCA, disruption of which abolished Gag binding to SeCTCA but not other tRNAs in cells. Importantly, the disruption of the MA-SeCTCA interface also diminished infectivity of progeny virions and impaired virus spread among CD4+ T cell lines. Altogether, our findings revealed two different modes of MA-tRNA binding, which can generate two different Gag pools in the cytoplasm, and identified the MA interface distinguishing SeCTCA from other tRNAs as a molecular determinant for infectious HIV-1 particle formation and virus replication in T cells.

## RESULTS

### Most tRNAs bound to cytosolic Gag lacking NC are the tRNAs present relatively abundantly in cells except for SeCTCA

To identify the tRNAs bound to Gag via MA, we expressed in HEK293T cells Gag protein constructs lacking the NC domain (delNC Gag) using CMV promoter-driven and Rev-RRE-dependent expression plasmids and performed photoactivatable ribonucleoside-enhanced cross-linking and immunoprecipitation (PARCLIP)-RNA sequencing (*9*, *32*) with some modifications. The Gag constructs used in the PARCLIP experiments were fused with yellow fluorescent protein (YFP) so that it can be recovered by immunoprecipitation using anti-GFP/YFP nanobodies. As described in the introduction, membrane binding of Gag could dissociate some tRNAs from Gag. Therefore, in addition to the delNC Gag that contains the WT MA sequence, we used a delNC Gag construct with a glycine-to-alanine substitution at the N terminus of MA, which blocks the Gag myristoylation (non-myr) and hence the attachment of Gag to the plasma membrane (*16*, *17*, *33*). To investigate the role of MA-HBR in tRNA binding, we also tested non-myr/6A2T delNC Gag in which basic residues of MA-HBR were substituted to neutral amino acids, threonine or alanine (Figure 1A) (*13*). Transfected cells were grown in a medium containing 4-thiouridine (4-SU), a photoactivatable ribonucleoside analogue, and then exposed to 365-nm UV light to crosslink 4-SU-containing RNA to the proteins they are bound to. Following cell lysis, delNC Gag proteins were immunoprecipitated, and RNA molecules bound to Gag were isolated by protease digestion of Gag protein followed by RNA extraction. Gag-bound RNA was run on a denaturing gel, and nucleic acids corresponding to size of tRNAs (60-100 nucleotides) were excised and recovered from the gel.

**Fig. 1:**
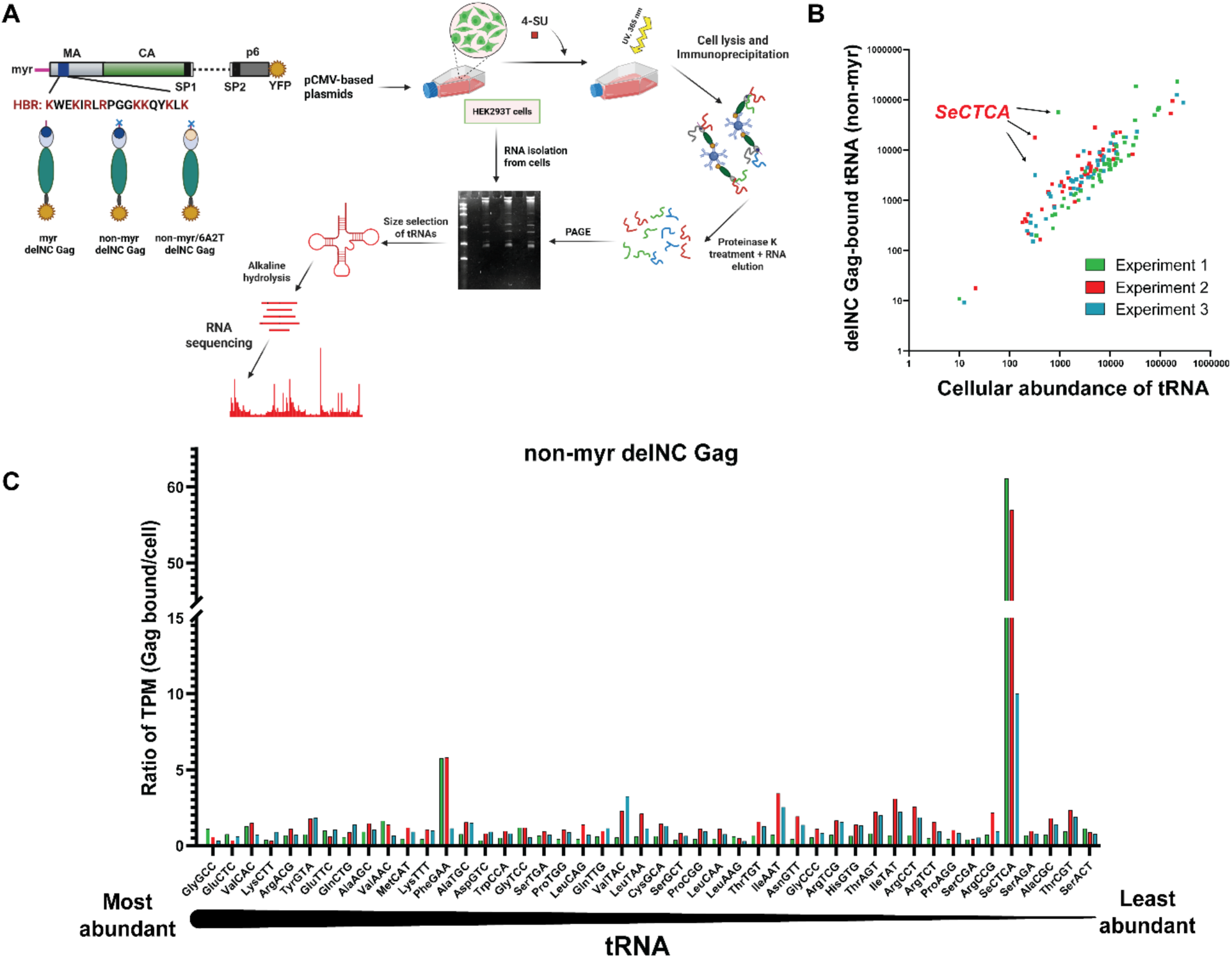
HIV Gag selectively binds Selenocysteine tRNA (SeCTCA) in cells. **(A)** Schematic illustration of the PARCLIP-hydro-tRNAseq. Myristoylation-deficient (non-myr), MA-HBR mutant (non-myr/6A2T), or myristoylated (myr) delNC Gag constructs, lacking most of the nucleocapsid domain and fused to YFP, were used as baits. Proteins were expressed in HEK293T cells, 4-thiouridine (4-SU) was added in the media, and RNAs containing 4-SU were crosslinked to their associated proteins with UV light. Subsequently, cell lysates were subjected to immunoprecipitation using an anti-GFP nanobody to isolate delNC Gag-YFP proteins, and RNA was dissociated from the recovered protein by proteinase K treatment. The recovered RNA was subjected to denaturing polyacrylamide gel electrophoresis, and RNAs with the size of tRNAs were extracted from the gel and sequenced following partial alkaline RNA hydrolysis. The histogram of the read length is shown in Figure S1A. **(B)** Number of reads corresponding to tRNAs present in non-myr delNC Gag-bound fraction versus tRNA abundance in HEK293T cells. Transcripts per million (TPM) values for total cellular and Gag-bound populations are plotted for each tRNA. In Experiment 1 (green), we plotted TPMs of Gag-bound tRNAs isolated from Gag-transfected cells to the TPM of total tRNAs isolated from cells transfected with pUC19, which represent tRNA population of generic transfected cells. In other two (Experiment 2, red; and Experiment 3, blue), we compared TPMs of Gag-bound tRNAs to the TPM of total tRNAs from cells expressing non-myr delNC Gag-YFP. The data for SeCTCA are indicated. The total raw read counts and the fractions of tRNAs bound to myr, non-myr, and non-myr/6A2T delNC Gag based on raw counts in the three experiments are shown in Figure S1B. **(C)** The ratios of TPM values for delNC Gag-bound tRNAs relative to TPM values of total cellular tRNAs isolated from HEK293T cells are shown. As described above, for the total cellular tRNA, we used the TPMs of tRNAs isolated from cells transfected with either a plasmid encoding pUC19 in Experiment 1 (shown in green) or non-myr delNC Gag-YFP in Experiments 2 and 3 (shown in red and blue). tRNAs are plotted on X-axis in the decreasing order of their cellular abundance based on the average TPM from the three experiments. SeCTCA enrichment was observed in case of non-myr delNC Gag-bound pool of tRNAs. See also Figure S2 for tRNAs bound to myr delNC and non-myr/6A2T delNC Gag constructs.

In this study, we coupled PARCLIP with the hydro-tRNAseq approach (*34*). Full-length tRNAs are difficult entities to reverse transcribe due to their thermodynamically stable secondary and tertiary structures and extensive base modifications (*35*). This inefficient reverse transcription (RT) might lead to low coverage and spurious measurements. To minimize the stalling of reverse transcriptase in cDNA synthesis step for maximizing tRNA read content in our tRNAseq analysis, we employed an RNA sequencing strategy based on partial alkaline RNA hydrolysis (hydro-tRNAseq), which generates shorter fragments of RNA with fewer secondary structure and base modifications (*34*). In this approach, the Gag-bound tRNAs derived from PARCLIP were subjected to alkaline hydrolysis and subsequently sequenced. To determine the relative abundance of tRNA species in HEK293T cells, we subjected total RNAs isolated from HEK293T cells to the same process as Gag-bound tRNAs. The reads from both Gag-bound and total cellular RNAs were mapped to a tRNA database curated by Gogakos et al. (*34*) based on human genome reference database (hg19; http://gtrnadb.ucsc.edu/). This tRNA database accounts for the sequence variations that occur upon reverse transcription of modified nucleotides by compiling edited reference transcripts with all possible combinations of modified nucleotide signatures at all detectably modified positions in the tRNA. This allows for mapping of nucleotide-modification-induced RT errors and hence enabling improved quantification of mature tRNAs.

An analysis of read counts for non-myr delNC Gag-bound tRNAs and cellular tRNA revealed that highly abundant tRNAs such as GlyGCC, GluCTC, ValCAC, or TyrGTA bound to Gag in relatively higher proportion in all our RNA sequencing datasets (Figure S2A). However, despite its low abundance in HEK293T cells, SeCTCA was overrepresented in non-myr delNC Gag-bound pool of tRNAs (Figure 1B and C). The comparison of Gag-bound versus total cellular tRNAs showed high enrichment of SeCTCA in all our sequencing datasets compared to other tRNAs including those that are abundant in cells. SeCTCA also showed higher enrichment in some of the datasets for myr delNC Gag-bound population of tRNAs (Figure S2B). In contrast, no consistent enrichment was observed for any of the cellular tRNAs with non-myr/6A2T delNC Gag (Figure S2C). Altogether, these results suggest that although most tRNAs bound to MA are those abundant in cells, SeCTCA, despite its paucity, binds MA efficiently presumably via interactions with basic amino acids in MA-HBR.

### MA binding to tRNA is dependent on MA-HBR and suppressed by MA binding to the PM

The PARCLIP-hydro-tRNAseq data presented above revealed the differences in tRNA populations that bind to Gag variants with different MA sequences, but due to the absence of control that allows for comparisons between different RNA pools in the PARCLIP-hydro-tRNAseq workflow, the magnitude of the effects of the MA amino acid substitutions on binding of each tRNA remained unclear. To address this, we aimed to compare the binding of the same tRNAs to different delNC Gag constructs (i.e., myr delNC Gag, non-myr delNC Gag, and non-myr/6A2T delNC Gag) using the PARCLIP followed by RT-qPCR. In this assay, we normalized bound tRNA levels between different Gag constructs at two steps. First, the amounts of the Gag proteins recovered after immunoprecipitation were quantified and used for normalization to account for differential transfection efficiency, protein expression levels, and immunoprecipitation yields. Second, the variability in RNA recovery and cDNA synthesis was normalized by spiking a 77-nucleotide long synthetic oligoribonucleotide (termed Spike RNA) in the samples before protease digestion of Gag-RNA complex recovered from PARCLIP (Figure S3). In the PARCLIP-RT-qPCR analysis, the spike RNA was quantitated along with tRNAs of interest using the corresponding primers (Table S1), and the yields of tRNAs are normalized by the spike RNA yields. For this analysis, we chose to quantitate a subset of tRNAs that are either detected abundantly in the delNC Gag-bound population or previously implicated in HIV-1 replication or binding to the Gag protein (Figures 1B and S2A). We found that the tested tRNAs displayed the highest binding to the non-myr delNC Gag variant among the three Gag variants (Figure S4A). In contrast, non-myr/6A2T delNC Gag showed tRNA binding similar to or below background levels for all the tested tRNAs, suggesting that the tRNAs bind to MA-HBR. For the analyses described thus far, Gag proteins were expressed using a CMV promoter-driven and Rev-RRE-dependent plasmid. To examine whether the expression of viral genomic RNA and other HIV-1 proteins would affect tRNA binding to Gag, we expressed the same variants of delNC Gag-YFP proteins in the context of a proviral molecular clone, pNL4-3 (Figure 2). As observed with the delNC Gag expressed by the CMV promoter-driven plasmid, delNC Gag encoded by pNL4-3 showed that MA-HBR is essential for tRNA binding to non-myr delNC Gag.

**Fig. 2:**
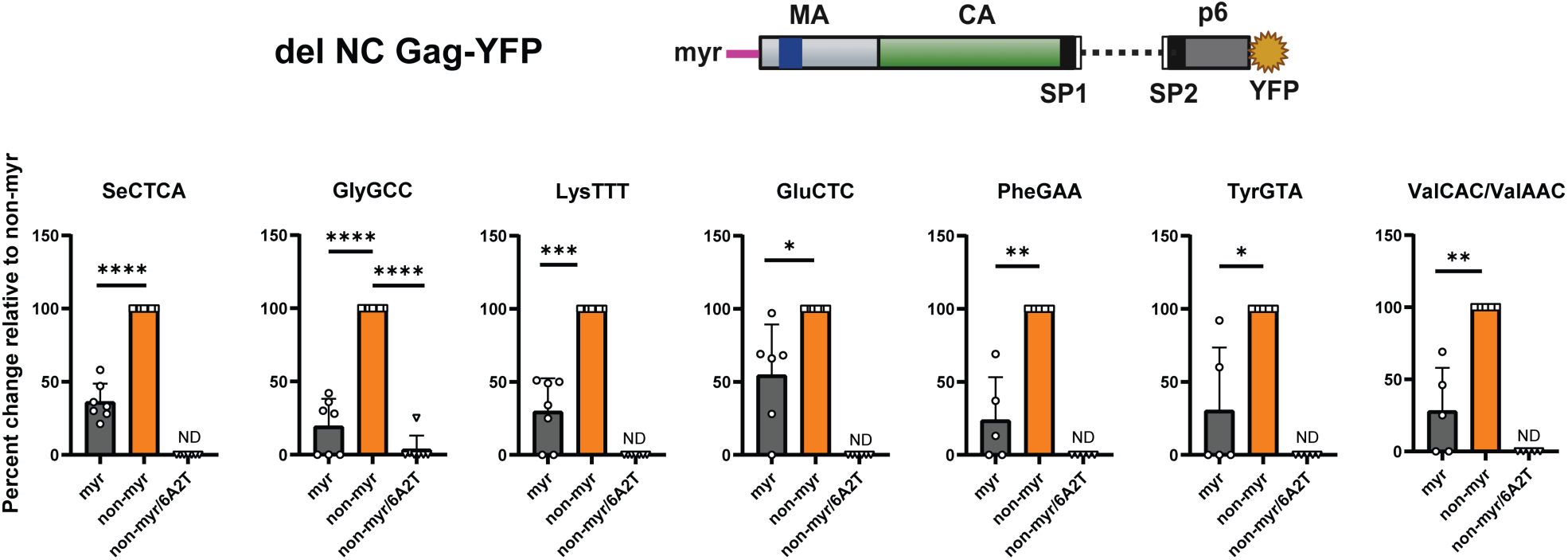
tRNA binding to delNC Gag is dependent on MA-HBR and partially reduced when Gag is competent in membrane binding. Transfection of HEK293T cells with plasmids encoding delNC Gag-YFP proteins or pUC19, photocrosslinking, and immunoprecipitation were performed as in Fig. 1. RNA was dissociated from the recovered protein by proteinase K treatment. A fraction of protein samples recovered after immunoprecipitation were run on SDS-PAGE and quantified after immunoblotting to measure relative levels of Gag proteins. An in vitro synthesized Spike RNA (Table S1) was added to the samples prior to RNA recovery to normalize for variations in RNA extraction and cDNA synthesis. cDNA was synthesized followed by protease digestion and subsequent phenol-chloroform extraction of RNA. A subset of tRNAs crosslinked to delNC Gag as well as Spike RNA were quantified using RT-qPCR as outlined in Figure S3. tRNAs bound to myr and non-myr/6A2T delNC Gag were compared with tRNAs bound to non-myr delNC Gag. Gag proteins were expressed using HIV-1 LTR promoter-driven plasmids. Protein levels and Spike RNA recovery were used for normalizing Cq values in RT-qPCR using specific primer pairs. ValCAC and ValAAC are indistinguishable with the primers used. Cq values higher or within 0.6 of the background, i.e., the values obtained with RNA recovered from pUC19-transfected (Gag-YFP-negative) cells using anti-GFP nanobody-bound magnetic beads, were assigned zero (See materials and methods). If data from all repeat experiments are zero for one condition, it is indicated as ND (not detected). The means ± SD of data obtained in seven independent experiments are shown. Of note, if the value for a given tRNA bound to non-myr delNC Gag is zero (when binding of a tRNA to non-myr delNC Gag did not reach the level higher than the background), we omitted the data set for that tRNA, since in this analysis we examined tRNA binding to the two other Gag constructs in comparison to the non-myr delNC Gag. Therefore, comparisons for some tRNAs were performed with fewer (but at least five) data sets than the rest of tRNAs. P value was determined using paired Student’s t test. *P < 0.05; **P < 0.01; ***P < 0.001, ****P < 0.0001. For non-significant comparisons, p values are shown.

The current working model for Gag localization at the plasma membrane entails that PIP2 in membranes outcompetes tRNA for binding to MA-HBR and facilitates specific association of Gag with the PM. Consistent with this model, we observed a significant reduction in binding of most tRNAs to myr delNC Gag, which is competent in membrane binding, compared to non-myr delNC Gag, especially when expressed in the context of the proviral clone (Figure 2). Altogether, these results indicate that tRNAs, including SeCTCA, GlyGCC, and LysTTT, bind Gag in an MA-HBR-dependent manner in the absence of NC and suggest that membrane binding of Gag displaces tRNAs from MA-HBR.

### SeCTCA binding to Gag is dependent on MA-HBR regardless of the presence or absence of NC

Next, we sought to determine the effect of the presence of the NC domain on tRNA binding to Gag. To this end, we expressed YFP-tagged full-length Gag constructs encoding different MA sequences (myr Gag, non-myr Gag, and non-myr/6A2T Gag). HEK293T cells were transfected with either CMV promoter-driven plasmid or HIV molecular clones encoding these Gag variants, and tRNAs bound to full-length Gag were analyzed by PARCLIP-RT-qPCR. Unlike delNC Gag, the full-length Gag with the non-myr/6A2T MA sequence showed similar or even higher binding to most tRNAs than those with non-myr or myr MA sequences, suggesting that the NC domain bound these tRNAs and thereby diminished the HBR dependence of Gag binding to these tRNAs (Figures 3 and S4B). Binding of ValCAC/ValAAC to full-length Gag was lower when MA HBR is disrupted but clearly detectable unlike with delNC Gag (Figures 2 and S4A), again suggesting the NC binding to the tRNA. In contrast, binding of SeCTCA to the full-length Gag variant bearing the non-myr/6A2T MA sequence was minimal compared to its binding to non-myr full-length Gag, indicating that binding to SeCTCA is strictly dependent on MA-HBR and that unlike other tRNAs, it does not bind the NC domain. The presence of the myristylation signal either significantly reduced or tended to reduce binding of all tested tRNAs to full-length Gag as observed for delNC Gag.

**Fig. 3:**
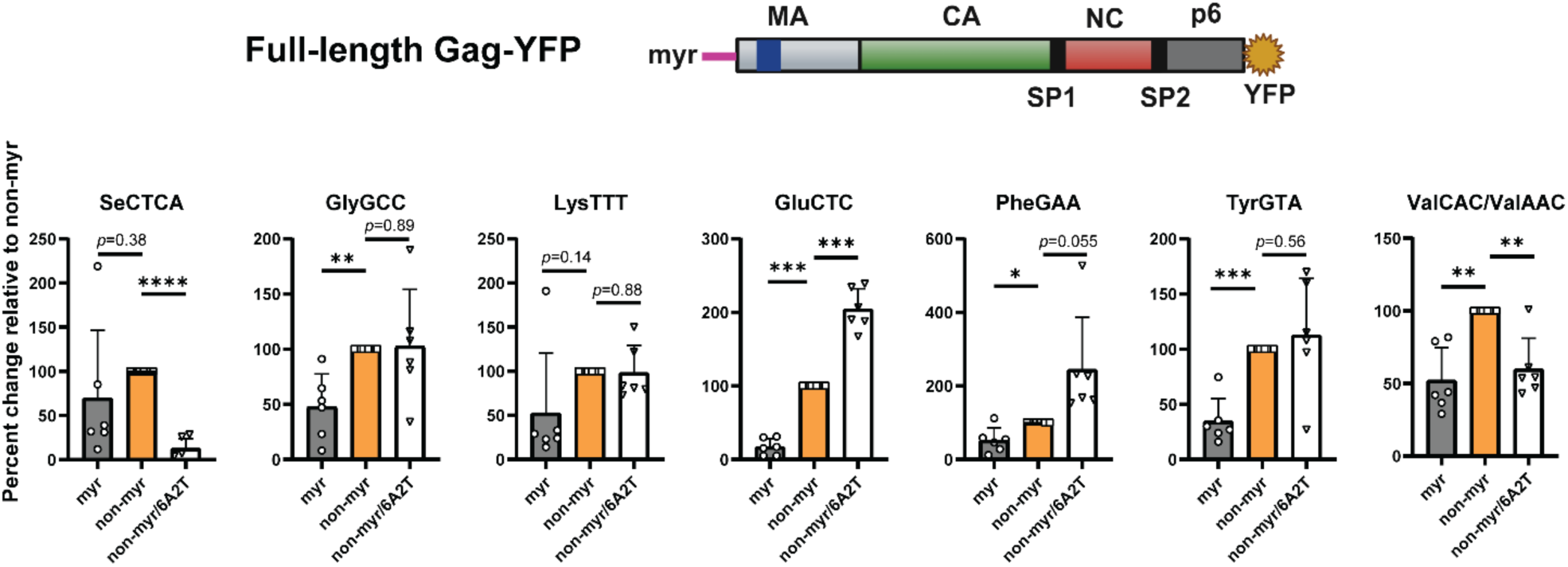
Binding of SeCTCA to full-length Gag is dependent on MA-HBR, whereas that of other tRNAs is not. A subset of tRNAs crosslinked to full-length Gag were quantified using PARCLIP-RT-qPCR as described in Fig. 2. tRNAs bound to CMV promoter-driven (A) or HIV-1 LTR promoter-driven (B) myr and non-myr/6A2T full-length Gag were compared with tRNAs bound to non-myr Gag. Protein levels and Spike RNA recovery were used for normalizing Cq values in RT-qPCR. Cq values higher than or within 0.6 of the background were assigned zero. The means ± SD of six independent experiments are shown. P value was determined using paired Student’s t test. *P < 0.05; **P < 0.01; ***P < 0.001, ****P < 0.0001. For non-significant comparisons, p values are shown.

### NC-mediated tRNA binding likely masks the requirement for HBR in tRNA-Gag binding

The analyses above revealed a considerable difference caused by the presence of the NC domain in the HBR dependence of Gag binding for non-SeCTCA tRNAs. It is possible that the direct binding of these tRNAs to NC masks the effect of HBR substitutions in MA. Alternatively, a change in availability of the MA-tRNA interface triggered by NC-dependent multimerization may explain why non-myr MA failed to show a higher tRNA binding than non-myr/6A2T MA in the context of full-length Gag unlike delNC Gag. To distinguish these two possibilities, we replaced the NC domain with a leucine zipper (LZ) dimerization motif in full-length Gag constructs. The use of GagLZ eliminates the effect of NC-mediated RNA binding, while promoting Gag multimerization even more efficiently than the NC domain (*36–39*). We examined the binding of different tRNAs to a panel of the GagLZ-YFP constructs containing myr MA, non-myr MA, and non-myr/6A2T MA using RT-qPCR. Unlike the observations with the full-length Gag constructs, all tested tRNAs showed significantly lower binding to the non-myr/6A2T GagLZ than to non-myr GagLZ (Figure S4C), suggesting that higher-order multimerization of Gag on the PM is not the reason for the loss of HBR-dependence of Gag binding to tRNAs. Rather, the direct tRNA binding to the NC domain, which may or may not be selective, likely accounts for the apparent loss of HBR-dependence of tRNA binding in the context of full-length Gag. Unlike the cases with delNC and full-length Gag, SeCTCA did not show a significant reduction in binding to myr GagLZ versus non-myr GagLZ (compare Figures S4C with S4A and S4B). These results suggest that SeCTCA, unlike other tRNAs, remains bound to GagLZ even when it is competent in membrane binding, possibly reflecting its efficient MA binding relative to other tRNAs.

### SeCTCA binding to MA suppresses interaction between Gag and membranes

Since GlyGCC, LysTTT, and SeCTCA bind to MA in an HBR-dependent manner in cells, we also examined whether these tRNAs can suppress MA-mediated membrane binding. To examine the effect of specific tRNAs on Gag interaction with non-PIP2 acidic phospholipids, we performed liposome binding assay using a purified recombinant myristoylated MA-CA-SP1 (myrMACASP1) protein and in vitro transcribed tRNAs (Figure 4A-C). In a similar assay using purified myristoylated MA protein (myrMA), we previously observed that LysTTT is capable of inhibiting myrMA-liposome binding (*28*). All the tested tRNAs showed strong inhibition of myrMACASP1 protein binding to liposomes containing neutral phosphatidylcholine (PC) and acidic phosphatidylserine (PS) at a 2:1 ratio (Figure 4D). SeCTCA tRNAs showed significantly higher inhibition of liposome binding than LysTTT. When a physiologically relevant concentration of PIP2 (2%)(*40*, *41*) was included in the liposomes, the inhibitory effect of all three tRNAs on liposome binding was reduced, confirming that PIP2 counteracts the tRNA-mediated inhibition of MA-membrane interaction (Figure 4E). Under this condition, SeCTCA still showed higher inhibition of liposome binding, suggesting a stronger binding of SeCTCA to the myrMACASP1 protein.

**Fig. 4:**
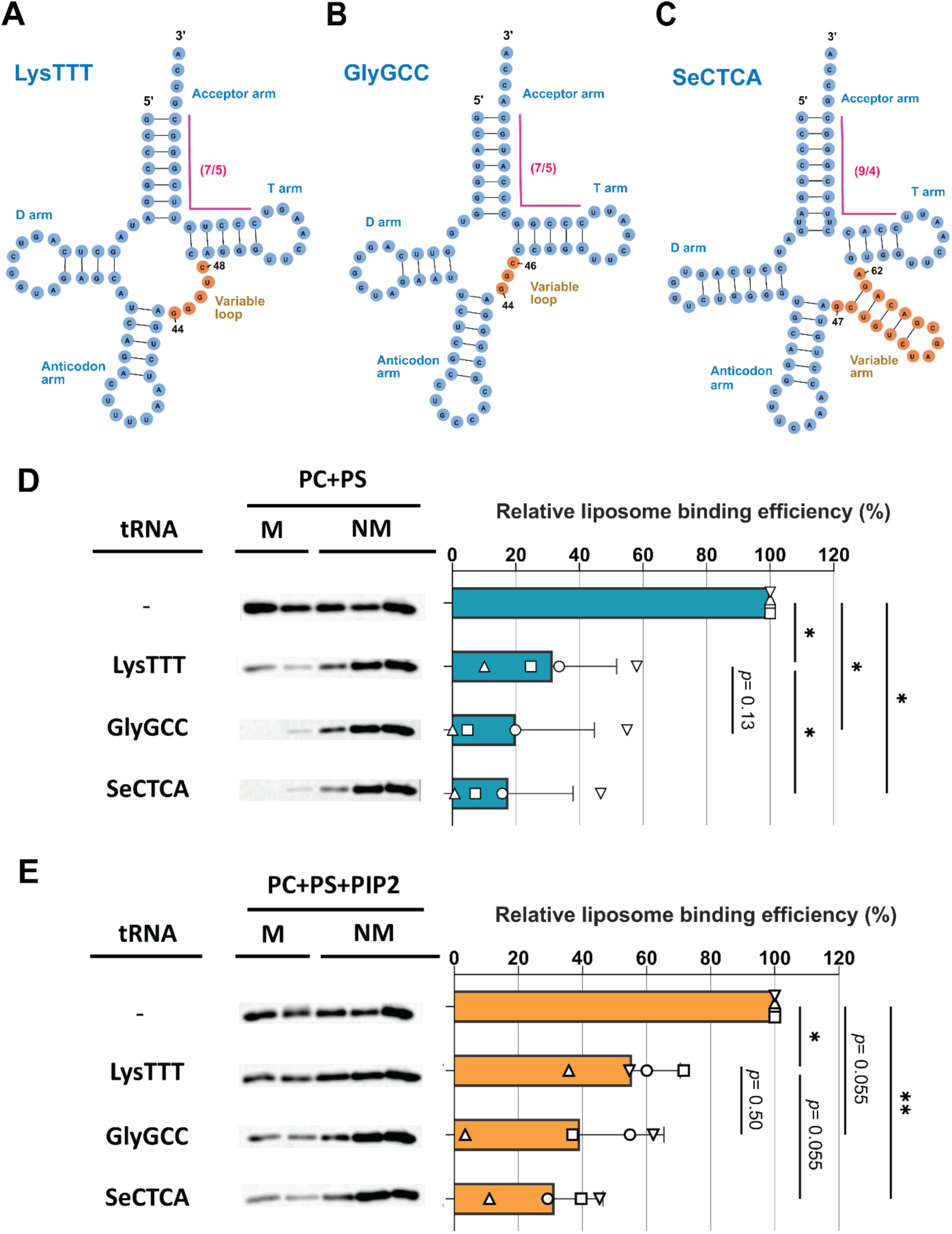
SeCTCA inhibits membrane binding of myristoylated MA-CA-SP1 (myrMACASP1) protein efficiently. **(A-C)** Two-dimensional representation of LysTTT (A), GlyGCC (B), and SeCTCA (C). SeCTCA has 9 and 4 bps in the stems of the acceptor arm and T arm, respectively, which give it the non-canonical 9/4 configuration, and a long variable arm (shown in orange). **(D-E)** tRNAs were in-vitro transcribed and incubated with the purified myrMACASP1strep protein. PC+PS liposomes (D) or PC+PS+PIP2 liposomes (E) were then added to each reaction, incubated for 1 h and subjected to equilibrium flotation centrifugation. Fractions were analyzed by SDS-PAGE followed by Western blot analysis of the protein with an anti-StrepTag antibody. PC, phosphatidylcholine; PS, phosphatidylserine; PIP2, phosphatidylinositol 4,5-bisphosphate; M, membrane-bound fractions; NM, nonmembrane-bound fractions. The relative liposome binding efficiency was calculated as the amount of membrane-bound Gag as a fraction of total Gag for each condition and was normalized by the liposome binding efficiency for the condition where no tRNA was added within the same experiment. The means ± SD of four independent experiments are shown. Data points of different replicates are indicated with distinct symbols (circle, square, and triangles). P value was determined using analysis of variance (ANOVA) and one-way Tukey’s multiple comparison test. *P < 0.05; **P < 0.01. For non-significant comparisons, p values are shown.

### MA residues Arg19 and Lys25 interact with the SeCTCA V arm, thereby forming an interface unique to the MA-SeCTCA complex

To further delineate the specific binding of MA to SeCTCA versus other tRNAs, we employed AlphaFold 3 to generate the structural models of the MA-SeCTCA and MA-GlyGCC complexes (Figure S5A and B)(*42*). A comparison between the predicted structure of the MA-SeCTCA complex and the previously determined crystal structure of a MA-tRNA complex wherein the tRNA was a modified Lysine tRNA (PDB ID: 7MRL)(*29*) revealed that MA binds to SeCTCA not only via interactions with its D- and T-arms as observed in the MA-tRNA crystal structure but also via contacts with the characteristic V arm of SeCTCA (Figure 5A-D). The loop connecting α1 and α2 helices of MA forms extensive contacts with SeCTCA, in a binding cavity lined by the protruding V-arm of SeCTCA on one side and D- and T-arms on the opposite (Figure 5B and C). Similar to the crystal structure of the MA-tRNA_Lys_ complex, AlphaFold-predicted models of both MA-SeCTCA and MA-GlyGCC complexes showed MA interactions with D- and T-arms via Arg21, Lys26, Lys31, His32, Trp35, Glu72, and Ser76 (Figures 5D and S5D). Additionally, in the MA-SeCTCA complex, two conserved residues of MA-HBR, Arg19 and Lys25, located in the α1-α2 loop, make extensive hydrogen bonds with the V-arm and T-loop of SeCTCA (compare Figure 5D with Figure S5C and D); Arg19 binds to the backbone oxygens of tA60 and tG61, and Lys25 forms a hydrogen bond with the phosphate oxygen of tU64.

**Fig. 5:**
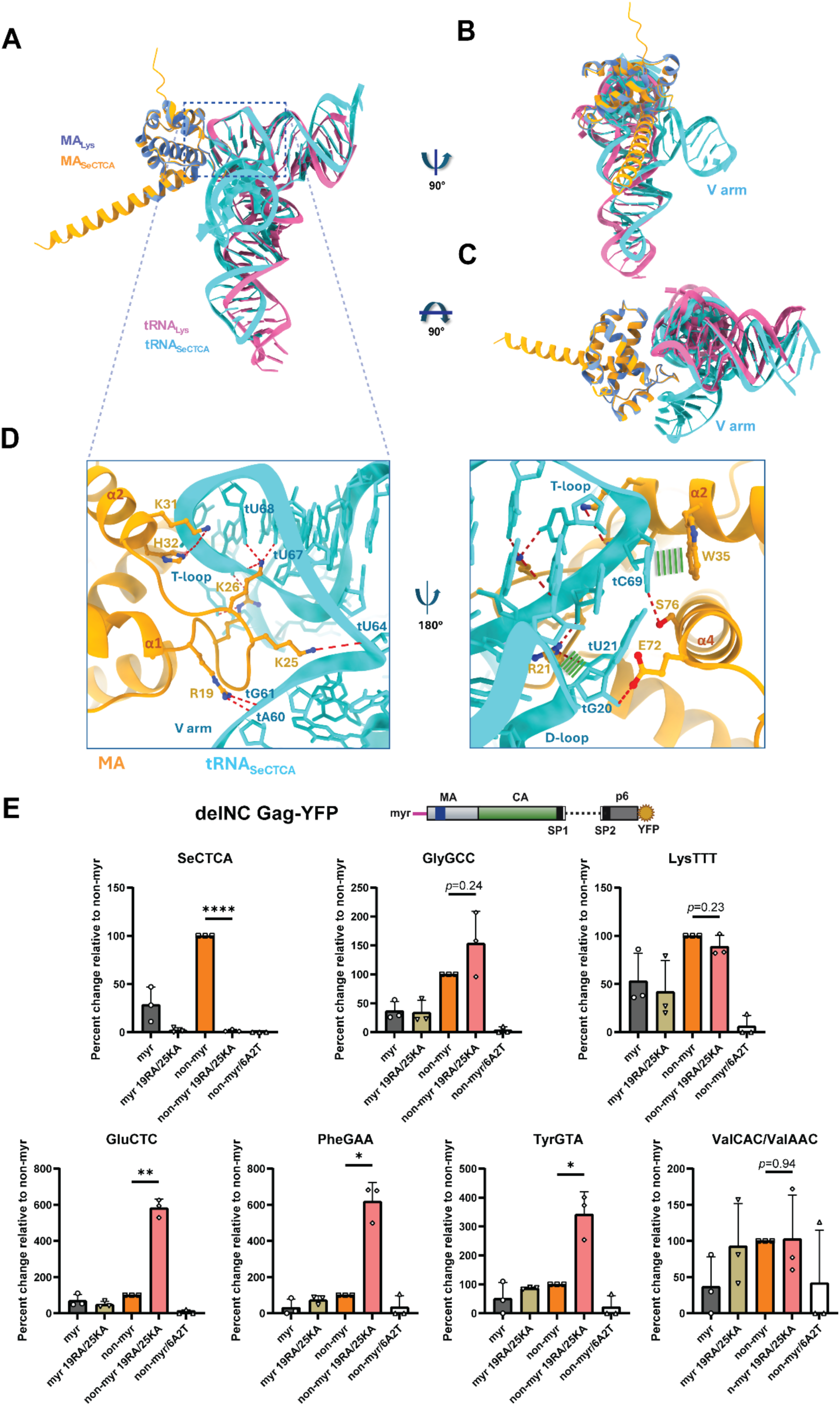
MA amino acid residues Arg19 and Lys25 in MA-HBR are the major determinants of SeCTCA binding to MA. The structure of the MA-SeCTCA complex was predicted using AlphaFold 3 (see also Figure S5). **(A)** Cartoon representation of the superposition of the AlphaFold 3-predicted MA-SeCTCA complex structure and the crystal structure of the previously determined complex between MA and a modified tRNALys (PDB ID:7mrl). The predicted local distance difference test (pLDDT) score of MA-SeCTCA complex model is shown in Fig. S5A. **(B-C)** Side (B) and top (C) views showing the elongated V arm of SeCTCA constituting an additional putative interface for MA. MA_SeCTCA_ and MA_Lys_ are shown in orange and blue, respectively, while tRNA_SeCTCA_ and tRNA_Lys_ are shown in cyan and magenta, respectively. **(D)** Zoomed-in views of the interphase between MA and SeCTCA in the predicted complex structure. MA and SeCTCA are shown in orange and cyan, respectively. Hydrogen bonds are shown as dashed red lines; stacking interactions as parallel green lines. Nucleotides in tRNAs are numbered with a “t” prefix. **(E)** Binding of a subset of tRNAs to different delNC Gag-YFP proteins (myr, myr 19RA/25KA, non-myr, non-myr 19RA/25KA, and non-myr/6A2T) is assessed by PARCLIP-RT-qPCR as described in Fig. 2. The means ± SD of three independent experiments are shown. P value was determined using paired Student’s t test. Only statistical significances of differences between non-myr delNC Gag and the corresponding 19RA/25KA mutant are shown for clarity. *P < 0.05; **P < 0.01, ****P < 0.0001. For non-significant comparisons, p values are shown.

In order to examine the role of the MA residues predicted to interact with SeCTCA V arm in binding of MA-SeCTCA interactions in cells, we introduced Ala substitutions at MA residues 19 and 25 (19RA/25KA) in delNC Gag-YFP and performed PARCLIP-RT-qPCR assay as described above. These amino acid changes nearly abolished the binding of SeCTCA to both myr- and non-myr delNC Gag, indicating that Arg19 and Lys25 residues are the key molecular determinants of MA-SeCTCA binding in cells (Figure 5E). Interestingly, we observed a significant concomitant increase in the binding of other tRNAs including PheGAA, GluCTC, and TyrGTA to the 19RA/25KA variant, indicating that SeCTCA outcompetes these tRNAs for binding to WT MA-HBR. However, binding of GlyGCC and LysTTT tRNAs remain largely unaffected by these mutations, suggesting that the binding of these tRNAs to WT MA is already saturated or optimal at their cytosolic concentrations.

To validate the predicted role for the V arm in the MA interaction, we examined the SeCTCA derivatives with 10- and 12-nucleotide V arm deletions (delV_10_ and delV_12_, respectively) for the ability to block MA-mediated liposome binding (Figure 6A-B). The predicted structures of these deletion constructs are largely superimposable to the intact SeCTCA (Figure 6C). We found that these constructs are markedly less efficient in inhibiting binding of myrMACASP1 to PC+PS liposomes (Figure 6D), indicating that the V arm is a key part of the interface for SeCTCA interaction with MA.

**Fig. 6:**
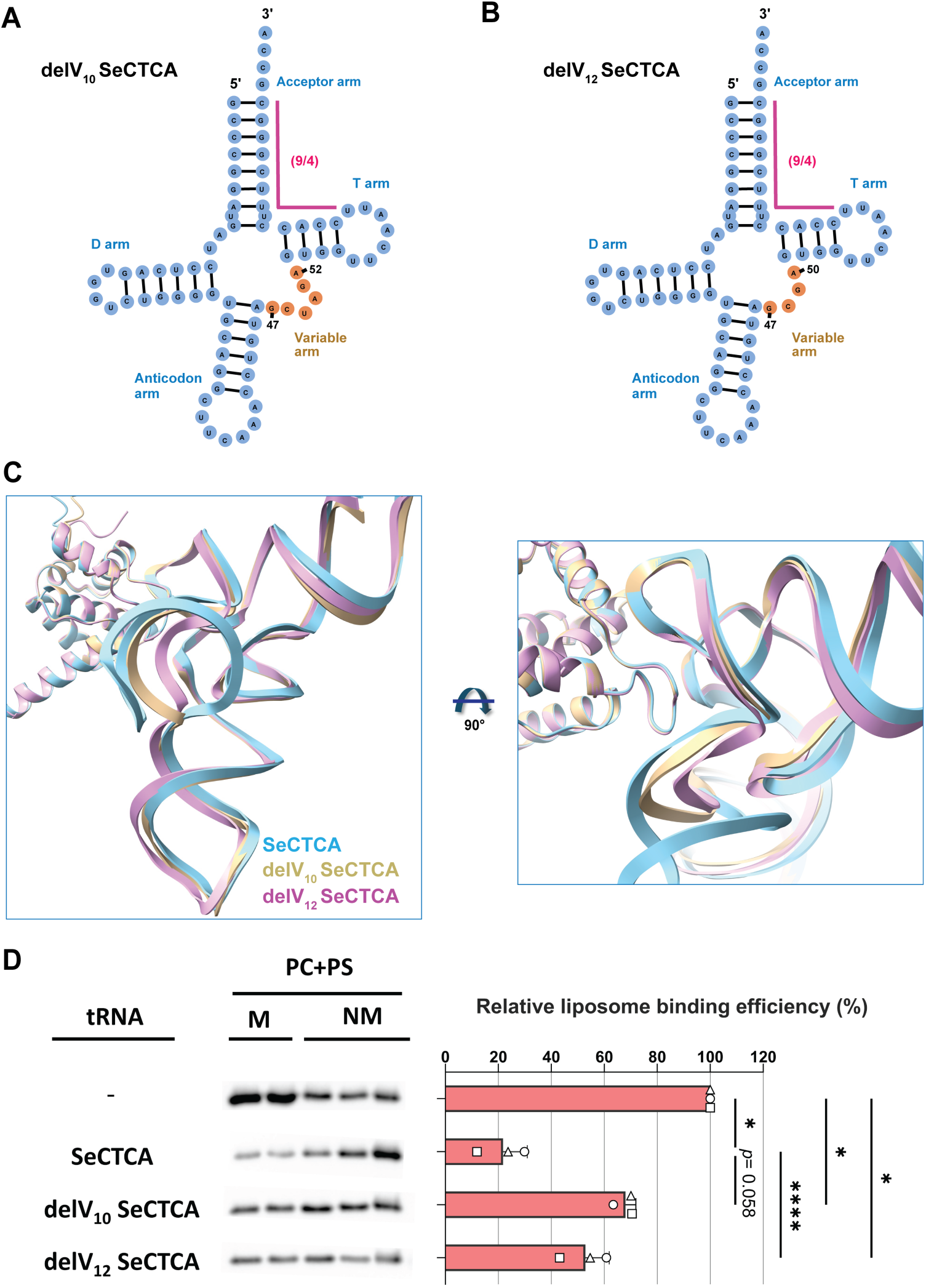
V arm is a determinant of SeCTCA-mediated inhibition of MA-dependent Gag-membrane binding. **(A and B)** Two-dimensional representation of delV_10_ SeCTCA (A) and delV_12_ SeCTCA (B) showing shortened V arms. **(C)** Front and top view of superposition of predicted model structures of HIV-1 MA complexed with SeCTCA, delV_10_ SeCTCA, or delV_12_ SeCTCA (shown in blue, yellow, and pink, respectively). Backbone of tRNA is shown for clarity. Note that overall conformations predicted for delV_10_ SeCTCA and delV_12_ SeCTCA are similar to that of SeCTCA. **(D)** Relative PC+PS liposome binding efficiency of myrMACASP1 Gag in the absence or presence of the tRNAs. The indicated tRNA or tRNA derivatives were in-vitro transcribed and incubated with the purified myrMACASP1strep protein. PC+PS liposomes were then added to each mixture, incubated for 1 h and subjected to equilibrium flotation centrifugation. Fractions were analyzed by SDS-PAGE followed by Western blot analysis of the protein with an anti-StrepTag antibody. PC, phosphatidylcholine; PS, phosphatidylserine; M, membrane-bound fractions; NM, nonmembrane-bound fractions. The relative liposome binding efficiency was calculated as the amount of membrane-bound Gag as a fraction of total Gag for each condition and was normalized by the liposome binding efficiency for the condition where no tRNA was added within the same experiment. The means ± SD of three independent experiments are shown. Data points of different replicates are indicated with distinct symbols (circle, square, and triangle). P value was determined using analysis of variance (ANOVA) and one-way Tukey’s multiple comparison test. *P < 0.05; ****P < 0.0001. For non-significant comparisons, p values are shown.

### Disruption of the MA interface for SeCTCA alters intracellular Gag localization and impairs infectivity of released virus particles

The loss of binding of SeCTCA prompted us to investigate the effect of the 19RA/25KA substitutions on viral replication. To this end, we introduced the 19RA/25KA changes in the pNL4-3 and pNL4-3/Gag-YFP and compared the MA variant with WT for Gag-YFP localization in HeLa cells and the efficiency of virion release from the HEK293T cells. We observed that 19RA/25KA Gag shows a reduced hazy cytosolic signal and increased cytoplasmic puncta relative to WT at 14 hours post-transfection (Figure 7A). However, Western Blot analysis of the cell-associated Gag and Gag released in virions showed no significant differences in Gag expression and efficiency of virus release between WT and the 19RA/25KA Gag (Figure 7B). Notably, infectivity assay using an indicator cell line, TZM-bl cells, as well as single-cycle infection of Jurkat T cells with nanoluciferase-encoding viruses revealed that virions released by HEK293T and HeLa cells expressing the 19RA/25KA Gag variant have a significantly reduced relative infectivity than the WT counterparts (Figure 7C-D; Figure S6A-B). Consistent with the role of SeCTCA in virion infectivity, we observed a modest but significant reduction in relative infectivity of virions when produced by HEK293T cells in which SeCTCA levels are reduced ∼50% using a CRISPR/Cas9 system targeting the *SeCTCA* gene (Figure S6C-D). Single basic-to-neutral substitution at MA residue 25 (25KA) reduces virion infectivity substantially but not as much as the double substitutions 19RA/25KA (Figure S6E). Interestingly, the virus with a basic-to-basic substitution at MA residue 25 (25KR) did not show the WT-level infectivity (Figure S6E). This observation suggests that simply having a positive charge at MA residue 25 may be insufficient for the full infectivity, consistent with the high conservation of amino acid sequences in this region (Figure S6F).

**Fig. 7:**
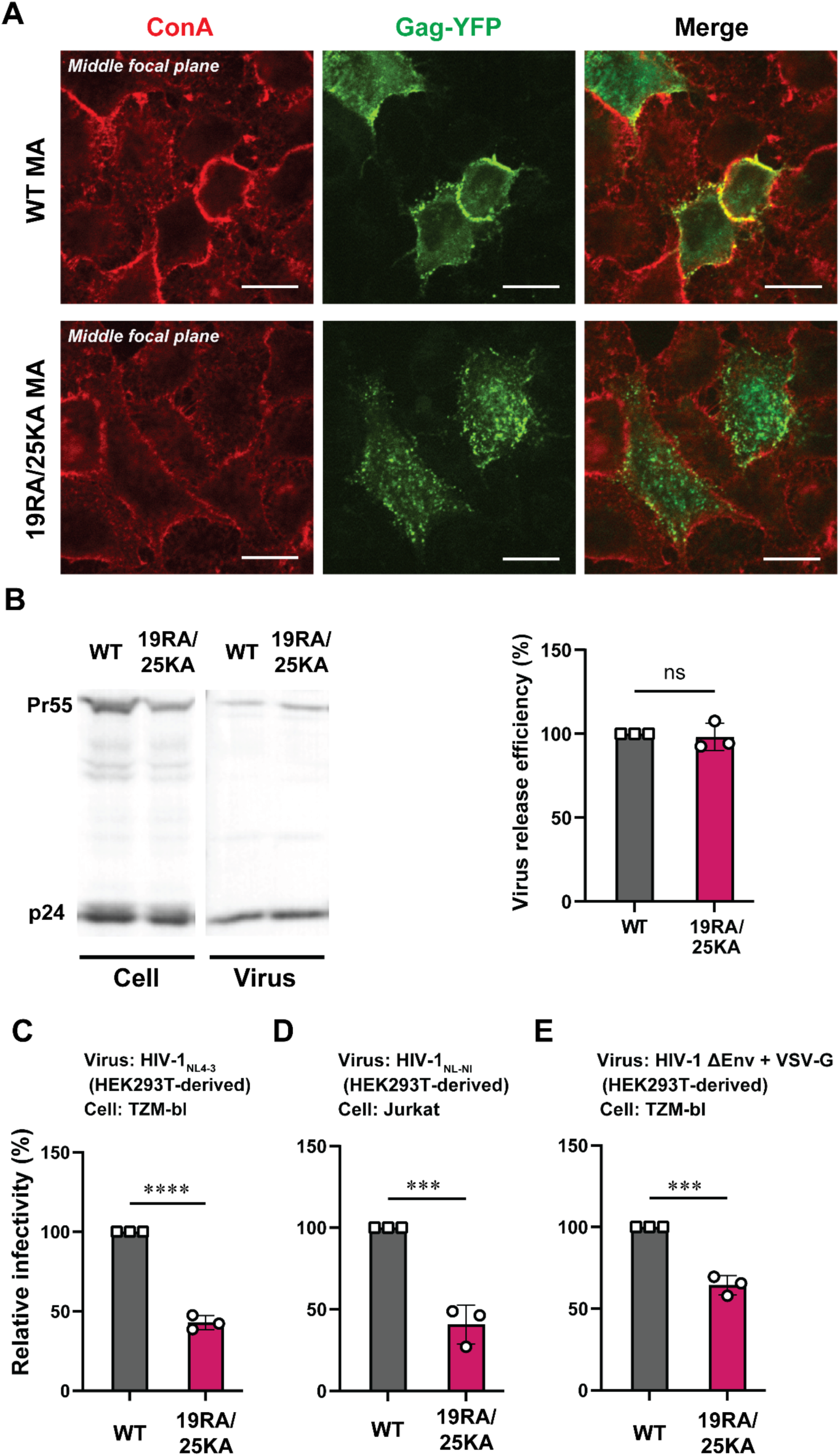
Loss of the MA interface for SeCTCA binding alters Gag localization in cells and impairs HIV-1 infectivity. **(A)** Representative images showing localization of Gag-YFP containing either WT or 19RA/25KA MA sequences. HeLa cells were stained with ConA-AF594 (red) at 14 h post-transfection and fixed with 4% paraformaldehyde in PBS. Localization of Gag (green) was analyzed using a confocal microscope. Middle focal planes of representative fields are shown. Scale bar, 20 µm. **(B)** HEK293T cells were transfected with WT or 19RA/25KA MA mutant. At 16 h post-transfection cell and virus lysate were collected and subjected to SDS-PAGE, and p24 and Pr55 proteins were detected by immunoblotting using HIV-Ig (left). Virus release efficiency of the MA mutant was calculated as a percentage of p24 released as pelletable materials in the supernatant to the total amounts of Gag (p24 released and p24+Pr55 present in cells) and compared to WT (right). The means ± SD of three independent experiments are shown. ns, not significant. **(C-D)** Single-cycle virion infectivity analysis was performed using TZM-bl cells (C) or Jurkat cells (D). For infectivity analysis in TZM-bl cells, HIV-1 was produced by transfecting HEK293T with pNL4-3 encoding WT or 19RA/25KA MA sequences. Sixteen h post-transfection, virus was collected and used to infect TZM-bl cells in the presence of saquinavir. At 48 h postinfection, cells were lysed, and luciferase activity was measured using the cell lysates. Luciferase activity was normalized by the amount of p24 in the inocula. For comparing virus infectivity in Jurkat cells, HEK293T were transfected with pNL-NI, a replication-competent HIV-1 molecular clone encoding NanoLuc and either WT or 19RA/25KA MA sequences, and the culture supernatant containing the virus was collected 16 h post-transfection. Jurkat cells were spinoculated with the virus at 25 °C for 2 h. Cells were then cultured for 24 h in the presence of saquinavir, and NanoLuc activity was measured in the cell lysate. NanoLuc activity was normalized by the amount of p24 in the inocula and total protein amount of Jurkat cells. **(E)** TZM-bl cells were infected with VSV-G-pseudotyped ΔEnv virus produced by cotransfection of HEK293T cells with pNL4-3/KFS and pHCMV-G. Luciferase activity was quantified and processed as in (C). All experiments were performed three times independently. Data are presented as mean ± SD. P value was determined using Student’s t test. ***P < 0.001, ****P < 0.0001.

To test whether the observed reduction in infectivity with the 19RA/25KA mutant is due to an Env incorporation defect, virion particles were generated by transfection of HEK293T cells with a VSV-G expression plasmid and pNL4-3/KFS (ΔEnv) encoding either WT or 19RA/25KA MA sequence. These VSV-G-pseudotyped particles showed a similar reduction in infectivity in the TZM-bl infectivity assay for 19RA/25KA MA as was observed with viruses that contain native Env, indicating that the effects of 19RA/25KA changes on virus infectivity is not accounted for by defects in Env incorporation or Env-mediated attachment (Figure 7E).

### Disruption of the MA-SeCTCA interface impairs spreading infection of HIV-1 in CD4+ T cell cultures

To address the significance of the MA-SeCTCA interface in HIV-1 replication in T cells, we examined virus spread in Jurkat and A3.01 T cell cultures that are transfected with molecular clones encoding WT or 19RA/25KA Gag (Figure 8A-B; Figure S7A-B). For a comparison, we also examined a molecular clone that encodes 31KA Gag, which was previously shown to be defective in binding to LysTTT and deficient in virus spread in some T cell lines (*29*). When compared to WT HIV-1, HIV-1 encoding 31KA Gag showed replication kinetics with only a 2-day delay in Jurkat cells and no or little delay in A3.01 cells in these experiments (Figure 8A-B; Figure S7A-B). In contrast, HIV-1 with the 19RA/25KA substitutions spread with 4-6 day delays in the peak of replication kinetics in Jurkat cells and either failed to spread at all or spread with ∼12-day delay in A3.01 cells. These results indicate that the MA interface for SeCTCA is essential for efficient virus replication in the T cell cultures in contrast to the previously identified interface for other tRNA(s). The delay in virus replication caused by the 19RA/25KA changes was also prominent when the T cell cultures were inoculated with HIV-1 wherein two-fold more inoculum was used for the 19RA/25KA virus than for WT virus to achieve the equivalent first-cycle infection (Figure S7C-D).

**Fig. 8:**
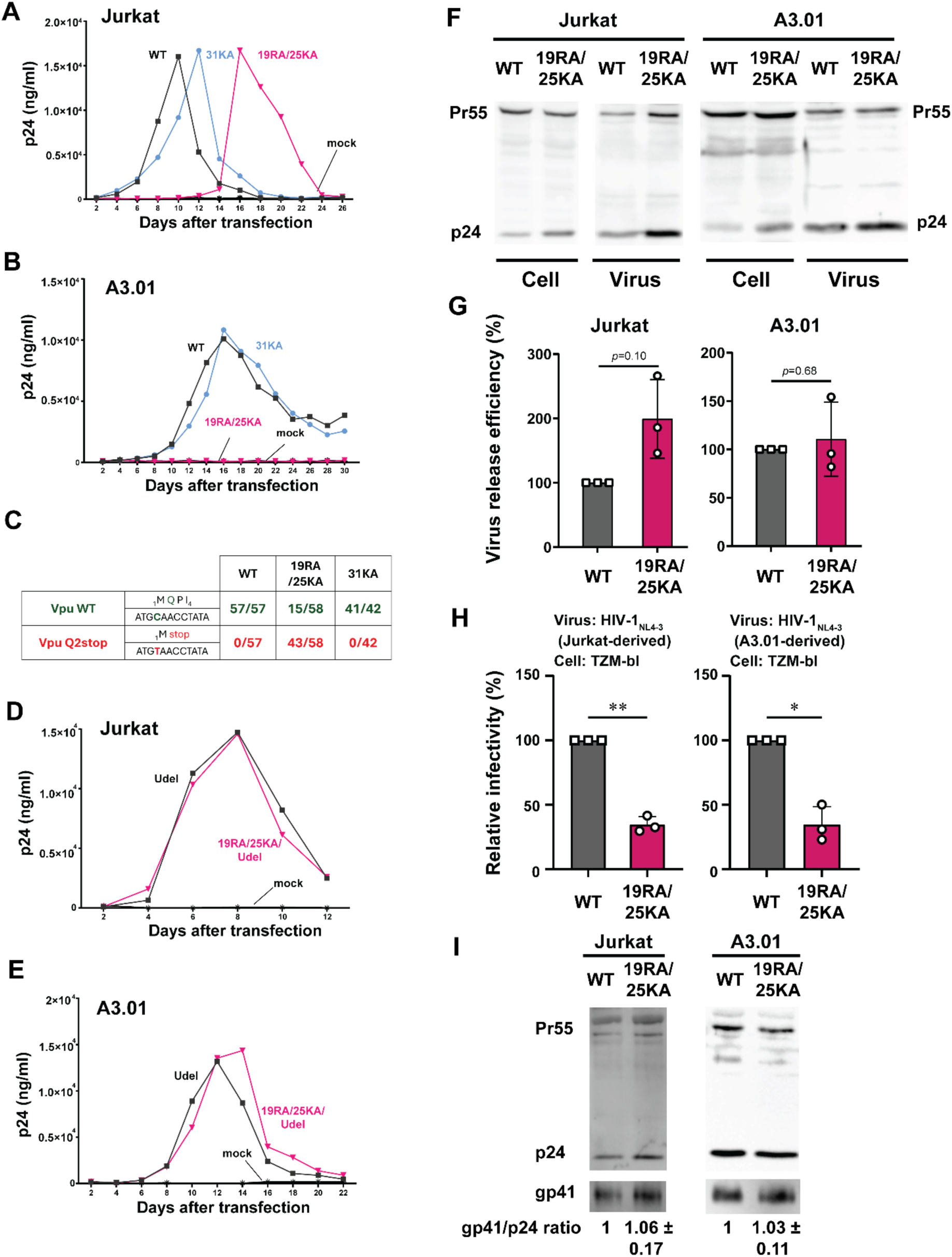
Loss of the MA interface for SeCTCA binding delays or abolish virus spread in CD4+ T cell line cultures. **(A-B)** Jurkat (A) and A3.01 (B) cultures were transfected with HIV-1 molecular clones that encode WT MA, 31KA MA, or 19RA/25KA MA using DEAE-dextran. Cells were split every 2 days, and virus spread was monitored based on virus released into the supernatants using p24 ELISA. One representative replication kinetics from two independent experiments is shown for each T cell line. A repeat experiment is shown in Fig. S7A-B. **(C)** Vpu substitutions in proviral sequences detected in the Jurkat culture. Frequencies of variant reads relative to total number of proviral sequence reads in sequencing of genomic DNA extracted at the peaks of replication in panel A are shown. **(D-E)** Jurkat (D) and A3.01 (E) cultures were transfected with Udel molecular clones that encode WT MA or 19RA/25KA MA using DEAE-dextran. Virus spread was monitored as in Panels A and B. **(F-G)** Jurkat and A3.01 cultures were inoculated with VSV-G-pseudotyped ΔEnv HIV-1 encoding either WT or 19RA/25KA Gag. Two-fold more inoculum based on p24 was used for the latter to compensate for the reduced infectivity associated with the 19RA/25KA changes. Cell-associated and virus-associated Gag were examined using immunoblotting (F), and virus release efficiency (G) was determined as in Figure 7B. **(H)** Jurkat and A3.01 cultures were inoculated with HIV-1 encoding intact Env and either WT or 19RA/25KA Gag. Two-fold more inoculum based on p24 was used for the latter to compensate for the reduced infectivity of the inoculum. The progeny virions released by the infected T cells were collected at 48 hours postinfection and examined for relative infectivity as in Figure 7C. All experiments were performed three times independently. **(I)** Virus-associated materials were examined for gp41 and Gag using immunoblotting. Relative Env incorporation efficiencies determined as the ratios of gp41 to p24 were compared between WT and the MA variant and shown at the bottom. Data are presented as mean ± variation from three (Jurkat) and two (A3.01) independent experiments. P value was determined using Student’s t test.*P < 0.05; **P < 0.01. For non-significant comparisons, p values are shown.

Notably, 74% of the proviral sequences obtained at the delayed peak of the 19RA/25KA virus (16 days postinfection) in Jurkat cultures from Figure 8A contain stop codon substitutions at the Vpu amino acid residue 2 (Figure 8C) while retaining the original 19RA/25KA changes. This is in a clear contrast to the proviral sequences at peaks of the WT and 31KA viruses (10 and 12 days postinfection, respectively) in Jurkat cultures where no *vpu* mutations were found. Disruption of Vpu expression enhances cell-to-cell spread (*43*, *44*), which is known to rescue virus replication in the conditions that would otherwise suppress virus replication. Therefore, these results suggest that the 19RA/25KA changes impose a selection pressure that is substantial enough to select for the variants with the *vpu* gene mutations. As expected, in the context of Vpu-deleted molecular clones (pNL4-3/Udel), the 19RA/25KA changes showed no or little effect on the virus replication kinetics in Jurkat and A3.01 T cells (Figure 8D-E).

To validate the phenotypes observed in HEK293T cells above (Figure 7) in the T cell lines, we infected Jurkat and A3.01 cells with VSV-G-pseudotyped ΔEnv HIV-1 encoding WT or 19RA/25KA Gag. For the latter, two-fold more inoculum by the p24 amount was used to compensate for the reduced infectivity (Figure 7C-E). We found no major changes in Gag expression and particle release in A3.01 cells (Figure 8F-G) although in Jurkat cells, virus particle release was enhanced by 19RA/25KA (Figure 8F-G). This may explain the observation that the impact of the substitutions on virus replication kinetics was less severe in Jurkat cells than in A3.01 cells (Figures 8A-B and S7). We then infected Jurkat and A3.01 cells with HIV-1 encoding intact Env and either WT or 19RA/25KA Gag (again, latter with two-fold more inoculum) and examined the relative infectivity of progeny virions collected 48 hours postinfection. We found that the defect in relative virion infectivity associated with 19RA/25KA Gag was slightly more severe with viruses produced in both Jurkat and A3.01 cells than with HEK293T-derived viruses (Figure 8H). Finally, we examined the efficiency of Env incorporation into virions produced by the T cells. We found that in both T cell lines, Env incorporation was unaffected by the 19RA/25KA substitution (Figure 8I). Altogether, disruption of the MA-SeCTCA interface does not cause defects in virus particle assembly or Env incorporation yet significantly suppresses virion infectivity and virus replication in CD4+ T cells.

## DISCUSSION

The earlier PARCLIP study that profiled the tRNAs bound to the Gag protein in cells used Gag constructs competent in binding the PM (*9*). Under this condition, Gag membrane binding might lead to the dissociation of tRNA subpopulation bound to Gag in the cytoplasm (Figure S8A) and hence underestimation of Gag-bound tRNAs. The abundance of total tRNAs present in cells was also not examined, making it impossible to determine whether MA has preferences for specific tRNAs. To address these points, we compared tRNAs bound to a membrane-binding-incompetent Gag with total tRNAs in cells, while incorporating the recent developments in tRNA sequencing (*34*). Our PARCLIP-hydro-tRNAseq analysis revealed that tRNAs decoding Gly, Glu, Lys, and Val, which were previously identified as the tRNAs bound to MA most frequently (*9*), are for the most part tRNAs abundantly present in HEK293T cells. Therefore, it is likely that MA binding to a subset of tRNAs is primarily determined by the frequency of encounter between a negatively charged tRNA molecule and a positively charged MA-HBR. Interestingly, SeCTCA, which is present in far less abundance in cells, exhibited selective binding to non-myr Gag as well as myr Gag likely due to the unique binding interface identified in this study. Consistent with the broad applicability of our results, during the preparation of this manuscript, another group reported that SeCTCA binds MA in T cell lines using native RNA immunoprecipitation and sequencing (*45*).

SeCTCA, which decodes stop codon UGA in mRNAs to the 21st amino acid selenocysteine, has non-canonical structural and functional features compared to other cellular tRNAs. It is by far the largest tRNA in the eukaryotes (87 nucleotides in humans) with a 19-nucleotide long variable (V) arm that folds into a stem-loop secondary structure. SeCTCA uniquely adopts a 9/4 secondary structure (9 bp making the stem of the acceptor arm and 5 bp making the stem of the T arm) as opposed to the 7/5 structure seen for other tRNAs (Figure 4A-C). An AlphaFold model of the MA-SeCTCA complex shows a prediction that the long V arm of SeCTCA interacts with MA, thereby constituting an additional binding interface that is absent in other tRNAs (Figure 5A-D). The predicted interface was validated using Gag liposome binding assays; SeCTCA variants with the V arm deletions are much less efficient in blocking Gag-liposome binding (Figure 6). Upon disruption of the MA side of this interface, MA binding to SeCTCA was specifically inhibited, while binding to other tRNAs was either unchanged or even increased (Figure 5E). Moreover, this disruption resulted in diminished infectivity of the progeny virus particles (Figures 7C-E and 8H) and abolished or markedly delayed virus spread in CD4+ T cell lines (Figure 8A-B). Virus assembly and Env incorporation, the two major processes the MA domain is known to promote, are unaffected by the disruption of the MA-SeCTCA interface (Figure 8F, G, and I). Therefore, these results suggest a hitherto poorly studied role for MA in regulation of virion infectivity, either during particle assembly or after particle release, which requires the SeCTCA interaction. Importantly, disruption of the MA-SeCTCA interface selects for an adaptive mutation in the *vpu* gene, which was not observed in the WT virus culture (Figure 8C). Mutations that favor cell-to-cell transmission, including *vpu* gene disruption, are known to be selected when disadvantages in virus replication are imposed by various antiviral pressures. They include CA amino acid substitution, Nef deletion, and treatment with IN inhibitors, among others (*46–48*). Disruption of the MA-SeCTCA interface is likely to impose a disadvantage that is also severe enough to select for the similar adaptive mutations. These findings collectively support the notion that SeCTCA is a bona fide cofactor for HIV-1 replication.

For most tRNAs, non-myr full-length Gag showed higher binding than membrane-binding-competent full-length Gag (Figure 3). This tRNA dissociation from Gag upon membrane binding is in line with previous finding by Kutluay et al. wherein the authors observed a far lower tRNA read count in virions compared to the Gag-bound tRNA in cells. An in vitro study also showed that under conditions which promote the Gag conformation with the exposed myristoyl moiety, Gag binds tRNA with lower affinity than its myristoyl-sequestered conformation (*31*). Collectively, these observations strongly support that membrane binding of Gag disfavors tRNA binding to MA.

Of note, in the context of GagLZ, the presence or absence of the myristoyl moiety does not significantly affect Gag binding to SeCTCA (Figure S4C). In in vitro liposome binding experiments, SeCTCA inhibits membrane binding of myrMACASP1 proteins efficiently even when PIP2 is present (Figure 4). Therefore, it is conceivable that when a Gag multimer binds membrane, SeCTCA bound to a Gag molecule within the multimer may remain bound to Gag because of the relatively high affinity to MA-HBR while other tRNAs detach from their cognate Gag molecules. Previous studies showed that HIV-1 virions incorporate a limited number of tRNAs in addition to the HIV-1 reverse transcription primer tRNA, Lys tRNA (*49–52*). It is possible that the virion-incorporated non-Lys tRNAs, including SeCTCA (*52*), are the Gag-bound tRNAs that survived the displacement from HBR upon membrane binding of Gag at the assembly sites. Whether SeCTCA incorporated into virions in this manner plays a role in modulating virion infectivity is currently under investigation.

Analysis of tRNA binding to delNC Gag, full-length Gag, and GagLZ indicated that tRNAs except SeCTCA binds the NC domain in addition to MA-HBR. Consistent with this observation, previous PARCLIP and RNA-coIP studies showed that NC can bind tRNAs (*9*, *45*) although it remains to be determined whether NC-tRNA binding is tRNA sequence-specific or whether it is solely dependent on the tRNA abundance in cells. Gag in the cytoplasm adopts a compact conformation wherein MA is in close proximity to CA or NC domains, and Gag transitions to a more linear conformation during particle assembly (*53*, *54*). It is tempting to speculate that non-SeCTCA tRNAs bind MA and NC domains simultaneously in the cytoplasm, maintaining the Gag protein in a compact conformation. In this context, the Gag molecules that bind viral genomic RNA in the cytoplasm via the NC domain would limit tRNA binding only to the MA domain. These Gag molecules may have a higher probability of binding the plasma membrane, since PIP2 in the PM may readily displace MA-bound tRNA (Figure S8B). Consistent with this hypothesis, Carlson et al. observed that addition of RNA corresponding to the 5′ UTR of HIV, but not other RNAs of similar lengths, reverses tRNA-mediated suppression of membrane binding of recombinant Gag protein (*55*). Alternatively, removal of the tRNAs from the MA upon membrane binding could leave NC poised to bind genomic RNA and allow Gag to participate in the NC-RNA-interaction-mediated assembly (Figure S8C). This model is consistent with previous studies showing that viral RNAs dimerize and bind Gag on the PM (*56*, *57*). It is also possible that two subpopulations of Gag are present in the cells, depending on which tRNA Gag binds in the cytosol: SeCTCA-bound Gag molecules, which are likely to have unoccupied NC, and the group of Gag bound to non-SeCTCA tRNAs, which likely binds PM more easily. Since multimerization of Gag has been found essential for selective viral RNA binding and retention at the PM (*57–61*), the ratio of the two Gag populations, their propensity to participate in a given Gag multimer, and their relative contributions to membrane binding of the Gag multimer may modulate the fate of the Gag multimer in the assembly and packaging process. Of note, since viral RNA can mediate Gag multimerization in various subcellular locations (*62*), the initial interactions between the two Gag subpopulations could occur either on the PM or in the cytoplasm or both.

While SeCTCA binding to Gag generates a subset of cytoplasmic Gag that potentially differs from the rest of Gag, the sequestration of SeCTCA by Gag can also perturb translation of host selenoproteins. These proteins play a pivotal role in maintaining redox homeostasis in the cells. HIV infection has been reported to alter the redox status of the infected cells by inducing oxidative stress (*63*). Evidence suggests that this oxidative stress might play an important role in disease progression and clinical outcomes associated with HIV (*64*, *65*). Therefore, the sequestration of SeCTCA by Gag might have roles beyond virion assembly including the modulation of redox homeostasis in infected cells. However, the molecular mechanism(s) by which HIV infection alters the redox status remains to be determined. Future analysis of the MA-SeCTCA interaction and its interface may provide insights into the possibilities for not only suppressing infectious virus particle assembly but also restoring the redox status altered by infection using RNA-based therapeutics.

## METHODS

### Plasmids

pCMVNLGagPolRRE Gag-YFP, pCMVNLGagPolRRE delNC/Gag-YFP, and pCMVNL GagLZ-YFP were described previously (*24*, *26*). The non-myr version of these plasmids, pCMVNLGagPolRRE 1GA Gag-YFP, pCMVNLGagPolRRE 1GA delNC/Gag-YFP, and pCMVNL 1GA GagLZ-YFP were generated by PCR mutagenesis. The non-myr/6A2T mutants pCMVNLGagPolRRE/1GA/6A2T Gag-YFP, pCMVNLGagPolRRE/1GA/6A2T delNC/Gag-YFP, and pCMVNL 1GA/6A2T GagLZ-YFP were constructed by replacing the BssHII-SpeI region of pCMVNLGagPolRRE Gag-YFP, pCMVNLGagPolRRE delNC/Gag-YFP, and pCMVNL GagLZ-YFP, respectively, with the corresponding region of pNL4-3/1GA/6A2T (*66*). pCMV-Rev has been described elsewhere (*67*) (a kind gift from S. Venkatesan, National Institutes of Health).

The HIV-1 molecular clones pNL4-3/Gag-YFP, pNL4-3/delNC/Gag-YFP, and Vpu deletion mutant, pNL4-3/Udel were described previously (*27*, *68–70*). 1GA and 1GA/6A2T variants of pNL4-3/Gag-YFP were constructed from HIV-1 molecular clone pNL4-3 (*71*) by replacing its BssHII-SpeI region with that of pCMVNLGagPolRRE 1GA Gag-YFP and pCMVNLGagPolRRE 1GA/6A2T Gag-YFP, respectively. 1GA and 1GA/6A2T variants of pNL4-3/delNC/Gag-YFP were constructed using a similar approach.

For use in liposome binding assays, the MA-CA-SP1 region was cloned in a pET-11-based (*18*) plasmid co-expressing MA-StrepTag along with yNMT described previously (*28*). pNL4-3/KFS (env-) (*72*), VSV-G expression plasmid pHCMV-G (*73*), and a replication-competent HIV-1 molecular clone encoding NanoLuc, pNL-NI (*74*), were described previously. 19RA/25KA substitutions were introduced in pNL4-3/delNC/Gag-YFP, pNL4-3/1GA/delNC/Gag-YFP, pNL4-3, pNL4-3/Gag-YFP, pNL4-3/KFS (env-), pNL4-3/Udel, and pNL-NI, and 19RA, 19RK, 25KA, and 25KR substitutions were introduced in pNL4-3 using standard molecular biology procedures. Sequences of primers used for cloning in this study are available upon request.

### Cells and antibodies

HeLa cells, HEK293T cells, and TZM-bl cells were cultured and maintained in DMEM (Corning, USA) supplemented with 5% or 10% fetal bovine serum (FBS, Invitrogen) and 50 units/mL penicillin-streptomycin (Gibco) as described previously (*13*, *75*). Human CD4+ T cell lines, A3.01 cells and Jurkat cells, obtained from NIH AIDS Reagent Program were cultured in Roswell Park Memorial Institute (RPMI) medium (Invitrogen) supplemented with 10% FBS and 50 units/mL penicillin-streptomycin (Gibco) at 37°C. GFP-Trap® Magnetic Agarose nanobodies were purchased from ChromoTek (gtma-100). Anti-HIV immunoglobulin (HIV-Ig) was obtained from NIH AIDS Research and Reference Reagent Program. The antibodies against GFP and StrepTag were procured from Millipore Sigma (SAB4701015) and Abcam (ab76949), respectively. Horseradish peroxidase-conjugated secondary antibodies used for Western blotting analysis are: goat anti-human IgG (Invitrogen; 31410), goat anti-mouse (Millipore; AP310P), goat anti-rabbit (Invitrogen; G21234), rabbit anti-goat (Invitrogen; 31402).

### PARCLIP

For PARCLIP experiments, 2*10^5^ HEK293T cells/well were seeded in each well of 6 well plates in Dulbecco’s modified Eagle’s medium supplemented with 10% fetal bovine serum (Gibco) with penicillium and streptomycin (Gibco). Next day, cells were transfected with either a plasmid encoding one of the Gag-YFP proteins or pUC19. Twenty-Four h post-transfection, the medium was replaced with the fresh medium containing 200 µM 4-thiouridine (4-SU). The cells were cultured for 16 h and then irradiated with 365-nm UV light at 300 mJ/cm2 to cross-link RNA-binding proteins and 4-SU-containing RNA. The cells were subsequently lysed with NP-40 lysis buffer (50 mM HEPES buffer [pH 7.5] containing 150 mM KCl, 2 mM EDTA, 0.5% [v/v] NP-40 substitute, and 0.5 mM dithiothreitol [DTT] and supplemented with a protease inhibitor cocktail and recombinant RNasin [Promega, N2515]). The cell lysate was treated with 60 U/ml RNase-free DNase (Promega, M6101) and then incubated with GFP-Trap® Magnetic Agarose beads for 1 h on a rotating shaker at 4°C. The beads were washed three times each with IP-wash buffer (50 mM HEPES-KOH [pH 7.5], 300 mM KCl, 0.05% (v/v) NP-40 substitute, and 0.5 mM DTT), High-salt wash buffer (50 mM HEPES-KOH [pH 7.5], 500 mM KCl, 0.05% (v/v) NP-40 substitute, and 0.5 mM DTT), and PNK buffer (50 mM Tris-HCl [pH 7.5], 50 mM NaCl, 10 mM MgCl2). To determine the amount of Gag proteins affinity bound to GFP-trap beads by Western blotting, 10% of bead suspensions were collected prior to protease digestion and heated in SDS-PAGE buffer (50 mM Tris HCl [pH 6.8], 2 mM EDTA-NaOH, 10% [vol/vol] glycerol, 2% [v/v] SDS, 100 mM DTT, and 0.1% [w/v] bromophenol blue) for 5 min at 95°C. For Western blotting analysis, protein samples were run on 10% SDS-polyacrylamide gel and bands were transferred onto a polyvinylidene difluoride (PVDF) membrane. The membrane was blocked using SuperBlock in TBS (Thermo Fisher Scientific, PI37515). Gag-YFP-specific bands were detected by anti-GFP primary antibody and anti-rabbit secondary antibody (see “Antibodies” section). The chemiluminescent signal was detected using either West Pico or West Femto chemiluminescence substrate (Thermo Fisher Scientific, PI34580 and PI34696, respectively) and recorded on a GeneSys image acquisition system (Syngene).

For normalization of variability in RNA recovery and cDNA synthesis between different samples for the RT-qPCR analysis, equal volume of a 77-nucleotide ‘Spike RNA’ (synthesized by IDT, sequence listed in Table S1) was added to the remaining GFP-Trap bead suspensions. On-bead digestion of Gag protein was performed by heating the beads at 50°C for 30 min in proteinase K buffer (100 mM Tris-HCl [pH 7.5], 50 mM NaCl, 10 mM EDTA, and 2% SDS) supplemented with 2 mg/ml Proteinase K (Millipore Sigma, 3115828001). Gag-bound RNA was recovered by phenol/chloroform/IAA extraction (25:24:1, Invitrogen) followed by ethanol precipitation using GlycoBlue (Invitrogen, AM9515) as a co-precipitant.

For PARCLIP-hydro-tRNAseq, extracted RNA was run on a 10% denaturing acrylamide gel with 8 M urea and tRNA-specific bands (∼60-100 nucleotides) were excised. The RNA was recovered from gel by shaking the gel fragments in Tris-EDTA buffer with 0.25% SDS for 16 h at room temperature followed by phenol/chloroform/IAA extraction as described above.

### Hydro-tRNAseq

The RNA extracted from PARCLIP was subjected to limited alkaline hydrolysis. RNA was incubated with alkaline lysis buffer (1mM EDTA, 50mM Na2CO3 and 50mM NaHCO3 [pH 9.8]) at 60°C for 10 min. The partially hydrolyzed RNA was precipitated using ethanol. Library preparation using small RNA library prep kit (NEB; E7330S) using barcoded primers and RNA sequencing (on Illumina Mi Seq) was done by the University of Michigan Advanced Genomics Core. The quality of each sample was checked by FastQC (v0.11.7). Problematic reads were trimmed using Trim Galore (v0.6.4_dev), which is a wrapper for the Cutadapt (v2.8) (http://journal.embnet.org/index.php/embnetjournal/article/view/200/458) and FastQC (v0.11.7) with default parameters (e.g., minimum read length of 20 nucleotides). The trimmed reads were transformed using FASTX-Toolkit (v0.0.14) and customized script, then mapped using the scripts and reference database shared by the Tuschl Lab (*34*). Burrows-Wheeler aligner (BWA) was used to map sequencing reads against a curated and annotated database of mature tRNAs (*34*). Assignment of reads was done as described previously (*34*). Briefly, reads mapped to more than one location were mapped following a hierarchical procedure based on the cellular abundance of each RNA class wherein mature RNAs were prioritized. The BWA version used was (v0.7.17) and allows up to two mismatches with each round of mapping with increasing error tolerance (up to 2 mismatches) while retaining the mapping location with the lowest mismatch count. Notably, the curated database developed by the Tuschl lab (*34*) used for mapping accounts for all possible combinations of modified nucleotide signatures (i.e., base change) at all detectable modified positions in the tRNA, hence minimizing mismatch count. If a read is aligned equally well to N number of mapping locations, read count is split equally between them in order to avoid artificial read count inflation. Finally, all reads mapped to mature tRNA sequences with or without the signatures of nucleotide modifications were summed for quantification of a given tRNA isoacceptor. Transcripts per Million (TPM) is calculated by dividing the read counts of an RNA by the length of the RNA, then dividing by the sum of all RNAs in the sample. The number is further normalized by multiplying by 10^6. Bioinformatics analysis was performed using Perl, R, and Python scripts, that are available upon request.

### PARCLIP-RT-qPCR

For PARCLIP-RT-qPCR, extracted RNA was used to synthesize cDNA using SuperScript III First-Strand Synthesis kit (Invitrogen, 11752050). tRNAs were quantified with RT-qPCR using tRNA- and Spike RNA-specific primers (listed in Table S1) and SsoAdvanced Universal SYBR Green Supermix following manufacturer’s protocol (Bio-Rad, 1725271). Results were analyzed using Bio-Rad CFX Manager, and Cq values obtained for tRNA primer pairs were normalized with corresponding Cq values of Spike RNA. Cq values for tRNAs from Gag-expressing cells that are within 0.6 from the Cq values obtained from cells transfected with the pUC19 control were considered background. The fold changes in the normalized Cq values for tRNAs bound to Gag variants were calculated in comparison to the normalized Cq values for the same tRNAs bound to the non-myr Gag variants, which were considered as 100%. The values were subtracted from the pUC19 background levels and then corrected/normalized for the relative protein levels calculated by Western blotting analysis of the protein recovered after immunoprecipitation and shown as percentage of corresponding non-myr Gag variant.

### Expression and purification of recombinant myristoylated HIV-1 myrMACASP1

The myrMACASP1-strep was expressed from a plasmid that co-expresses the yeast N-terminal myristoyl transferase (yNMT) in E. coli BL21(DE3) pLysS. A primary culture grown overnight in LB supplemented with ampicillin (100 µg/mL) and chloramphenicol (34 µg/mL) at 37°C was used as an inoculum for a large-scale culture in Terrific Broth (TB) enriched with 4% glycerol, which was grown at 37°C to OD₆₀₀ 0.6–0.8. The cultures were cooled to 20°C and supplemented with myristic acid (10 µg/mL) 90 min prior to induction with 0.5 mM IPTG. Cells were collected 24 h later and were stored at −80°C until lysis. Cell pellets were resuspended on ice in Lysis Buffer (50 mM sodium phosphate pH 8.0, 500 mM NaCl, 10 mM DTT, cOmplete protease inhibitor tablets [Roche]) supplemented with 1 mg/mL each of Lysozyme (Thermo scientific; 89833) and DNase I (Thermo scientific; J62229.MC), and cells were lysed by sonication. The lysates were clarified by centrifugation at 35,000 g for 30 min at 4°C. To remove nucleic acids bound to proteins, 0.3% (w/v) Polyethylenimine (PEI) added to the cleared lysate, and the precipitated nucleic acids were removed by centrifugation at 20,000 g, 30 min, 4 °C. The resulting supernatant was filtered with 0.45-µm filter (Millipore; SLHVR33RB) incubated with pre-equilibrated Strep-Tactin®XT 4Flow® high-capacity resin (IBA Lifesciences; 25030002) for 15 min, collected by brief centrifugation (500 g, 1 min). The resins were washed three times with Buffer W (IBA Lifesciences; 21003100), and the protein with a Strep tag was eluted in Buffer BXT (IBA Lifesciences; 21042025). The protein eluate was buffer-exchanged to the assay buffer (100 mM Tris pH 7.5, 150 mM NaCl, 10 mM DTT) using 3 kDa MWCO Amicon Centrifugal Filter Device (Millipore) and stored at −80°C after purity of the protein was confirmed with SDS-PAGE.

### Liposome preparation

Liposome preparation was done as described previously with some modifications (*28*). Briefly, 1-Palmitoyl-2-oleoyl-sn-glycero-3-phosphocholine (POPC), 1-palmitoyl-2-oleoyl-sn-glycero-3-phospho-L-serine (sodium salt) (POPS), and L-α-phosphatidylinositol-4,5-bisphosphate (PIP2) (Avanti Polar Lipids Inc) in chloroform or chloroform:methanol:water = 20:9:1 were mixed at the indicated molar ratios (PC:PS = 2:1; or PC:PS:PIP₂ = 65.33:32.67:2.0) to a final total lipid concentration of 20 mM in the stock. Lipids were vortexed, dried under N₂ to a thin film, and placed under vacuum overnight. Films were resuspended in buffer with 20 mM HEPES, 50 mM NaCl (pH 7.5). The suspensions were subjected to 10 freeze–thaw cycles and were extruded through 1 µm polycarbonate membranes using a mini-extruder. Liposomes were made fresh for each experiment.

### Preparation of in vitro transcribed tRNAs

Human tRNAs (GlyGCC, LysTTT, and SeCTCA) were in vitro transcribed using T7 RNA polymerase (Promega, P2077). The primers used for amplification of tRNA genes to be used as templates for transcription are listed in Table S1. To suppress the non-specific N+1 activity of the T7 RNAP, primers were designed with two consecutive 2´-O-methyl modifications on 5´-end of the template strand. delV_12_ SeCTCA and delV_10_ SeCTCA were in vitro transcribed using annealed oligonucleotides (synthesized by IDT; listed in Table S1) using the same procedure. The in vitro transcribed tRNAs were purified by running the reaction mixture on 10% denaturing acrylamide-tris-borate-EDTA gel with 8 M urea. The tRNAs were extracted from the gel by phenol/chloroform/IAA extraction as described above. Recovered RNA was resuspended in water and refolded. For refolding, RNA was heated to 80°C for 2 min and incubated at 60°C for 2 min. Subsequently, RNA dissolved in water was adjusted to 1X refolding buffer (20 mM HEPES-KOH, pH 7.0, 100 mM KCl, 0.5 mM MgCl_2_) by addition of an appropriate amount of 10X refolding buffer and incubated at 37°C for 15 min. RNA was stored at 40 µM concentration at -80°C until further use.

### Liposome flotation assay (membrane binding)

Purified myrMACASP1-strep was incubated with in vitro transcribed tRNAs (10 µM each) in folding buffer (20 mM HEPES-KOH pH 7.0, 100 mM KCl, 0.5 mM MgCl₂) at 30°C for 1 h. This protein-tRNA mixture (6 µl) was incubated with 9 µl of extruded liposomes and incubated at 30°C for 1 h. 12.5 µL of the reaction were mixed with 87.5 µL of 40.3% (w/v) sucrose in 20 mM HEPES to a final 35.3% sucrose. Gradients were overlaid with 200 µL 25% and 40 µL 0% sucrose layers, then centrifuged at 67,600 × g, 2 h, 4°C. Four 80 µL fractions were collected from the top; any residual pellet material was resuspended in 60 µL 47% sucrose for analysis. Fractions were mixed with 2×SDS sample buffer, heated at 95°C for 10 min, separated by SDS-PAGE (10% acrylamide), transferred, and probed with anti-Strep-tag II primary (Abcam ab76949) and HRP-conjugated goat anti-rabbit secondary antibody. Chemiluminescent signals were recorded (West Pico Plus) and quantified. Liposome binding efficiency was calculated as the Gag signal in the top 2 fractions (membrane fractions) divided by the total Gag signal across all fractions.

### MA-tRNA complex modeling and structural analysis

The amino acid sequence of the HIV-1 MA protein derived from Genbank (ID: <u>AHN10731.1</u>) and tRNA sequences retrieved from genomic tRNA database (http://gtrnadb.ucsc.edu/; hg19) were used as inputs into the AlphaFold 3 server. The resulting coordinate file with highest confidence score was used for structural analysis using icn3d (*76*) and ChimeraX (*77*). pLDDT score visualization and superposition was done using ChimeraX.

### Virus production and viral release assay

To generate virus, 4*10^5^ HeLa or HEK293T cells were seeded in each well of 6 well plate and cultured for 24 h. Cells were transfected with WT or mutant MA-encoding plasmids, pNL4-3 and pNL-NI, to produce HIV-1_NL4-3_ and HIV-1_NL-NI_, respectively. To generate VSV-G-pseudotyped virus, HEK293T cells were co-transfected with pHCMV-G and either WT- or 19RA/25KA MA-encoding pNL4-3/KFS (ΔEnv). Transfection was performed using Lipofectamine2000 (Invitrogen) as described previously (*75*). At 16 h post-transfection, supernatants were filtered through a 0.45-μm filter. The filtered supernatants were used for p24 ELISA and virion infectivity assay. For analysis of virus release for HIV-1_NL4-3_, cell and virus lysates were prepared as previously described (*68*).

To examine virus release efficiency in infected T cells, 2*10^6^ cells were inoculated with 0.06 ng and 0.12 ng (in p24) HEK293T-derived VSV-G-pseudotyped ΔEnv HIV-1 (with WT and variant MA sequence, respectively) using spinoculation as described previously (*78*). To produce HIV-1_NL4-3_ in T cells, 2*10^6^ cells were inoculated with 2.5 µg and 5 µg p24 of 293T-derived HIV-1_NL4-3_ and its MA variant, respectively, by spinoculation followed by extensive washing. Forty-eight hours after spinoculation, culture supernatant and cell lysate were assayed for virus release, env incorporation or virion infectivity. Cell lysates were resolved on a 10% SDS-polyacrylamide gel, and immunoblotting was performed using HIV-Ig after transfer onto a PVDF membrane. Virus release was calculated using the following formula: 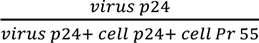

### Analysis of virus replication kinetics

To monitor viral relication in Jurkat and A3.01 cells, T cells were transfected with molecular clones based on pNL4-3 1 μg/1*10^6^ cells in the presence of 700 μg/mL DEAE-dextran (Millipore Sigma, 93556-1G) at 37°C for 15 minutes or were infected with 2.5 ng (in p24) HIV-1_NL4-3_ or 5 ng 19RA/25KA HIV-1_NL4-3_ using spinoculation. Transfected or infected cells were grown at 37°C, and cells were split 1:3 every other day with fresh medium and virus in culture supernatant was assayed using p24 ELISA. For analysis of viral genome, genomic DNA was extracted from cells using DNeasy Blood and Tissue Minikit (Qiagen) at the peak of viral replication. Viral DNA was amplified as two overlapping half-length amplicons (5’LTR-*vif* and *pol*-3’LTR) by PCR using PfuUltraII polymerase (Agilent Technologies, 600670) and primers (listed in Table S1). Amplicons were sequenced using Nanopore sequencing (Eurofins Genomics).

### p24 ELISA

p24 ELISA was done as described previously. Briefly, supernatants containing virus particles were lysed in lysis buffer (0.05% Tween 20, 0.5% Triton X-100, and 0.5% casein in 1XPBS). Nunc MaxiSorp plates were coated with an anti–HIV-1 p24 antibody (clone 183-H12-5C; NIH AIDS Research and Reference Reagent Program) and stored overnight at 4°C. Appropriate dilutions of lysed samples were added to the plate and incubated for 2 h. Captured p24 was detected calorimetrically with sequential incubation with biotinylated anti-HIV-1 p24 antibody (clone 31–90-25; American Type Culture Collection), streptavidin-poly HRP20 conjugate (Carbosynth), and 3,3′,5,5′-tetramethylbenzidine substrate (Sigma). Biotinylation of anti-HIV-1 p24 antibody was done using the EZ-Link Micro Sulfo-NHS-Biotinylation Kit (Pierce). p24 concentrations were determined using the curve generated with recombinant HIV-1 IIIB p24 protein as standards (NIH AIDS Reagent Program).

### Virus infectivity assay

To measure single-cycle virion infectivity of virus produced from HEK293T, HeLa, or T cells, 4*10^4^ TZM-bl cells were inoculated with 5-50 ng of virus (as determined by p24 ELISA) and cultured in the presence of 10 µM saquinavir (SQV; NIH AIDS Research Reagent Program). After 48 h of incubation, cells were lysed, and luciferase activity was measured with Luciferase Assay System (Promega, E1500) following manufacturer’s instructions. Luciferase activity was normalized by the amount of p24 in the inocula. To examine HIV-1 infectivity of virus collected from HEK293T or HeLa cells in T cell lines, 1*10^6^ Jurkat cells were infected with viruses produced by transfection of HeLa or HEK293T cells with pNL-NI or pNL-NI/19RA/25KA. Cells were further cultured at 37°C in the presence of 10 μM SQV. Cells were washed after 24 h and lysed with cell culture lysis reagent (Promega). NanoLuc activity was measured with Nano-Glo® Luciferase Assay System (Promega) following manufacturer’s instructions. The NanoLuc activity was normalized by p24 values (determined by p24 ELISA) and total protein amounts determined using the Pierce^TM^660 nm Protein Assay Reagent (Pierce).

### CRISPR-Cas9-mediated knockdown

To reduce SeCTCA level in the cells, single guide RNA (sgRNA) was designed and then cloned into the pSpCas9(BB)-2A-Puro (PX459) V2.0 (Addgene, #62988). HEK293T cells were transfected with the PX459 V2.0 plasmid carrying sgRNA sequence targeting *SeCTCA* gene sg-Forward (5’-CACC GAGTTACTACCGCCCGAAAGG-3’) and sg-Reverse (5’-AAAC CCTTTCGGGCGGTAGTAACTC-3’). PX459 V2.0 empty vector served as a control. Twenty-four hours post-transfection cells were selected with puromycin (4 µg/ml). After 5 days of selection, cells were plated in a new 6 well plate in puromycin-free DMEM containing 10% FBS and were transfected with pNL4-3 after 24 h. At 16 h post-transfection, supernatant was collected for p24 ELISA and virion infectivity assay. To analyze SeCTCA levels in the cells, total RNA was isolated from HEK293T cells using RNA miniprep kit (Zymo Research) and treated with RNase-free DNase (Promega, M6101), using manufacturer’s instructions. cDNA was synthesized, and RT-qPCR was performed as described in the PARCLIP-RT-qPCR section. Relative expression of SeCTCA was calculated by the 2^−ΔΔCt^ method using glyceraldehyde-3-phosphate dehydrogenase *(GAPDH)* as a control gene. Primers used for RT-qPCR included *SeCTCA*-specific primers listed in Table S1, and GAPDH-Forward (5’-ACCACAGTCCATGCCATCAC-3’) and GAPDH-Reverse (5’-TCCACCACCCTGTTGCTGTA-3’). The oligonucleotides were synthesized by IDT.

### Microscopy analysis

To determine the cellular localization of Gag, HeLa cells were plated onto an 8-well glass bottom chamber slide and cultured for 1 day at 37°C. Cells were transfected with pNL4-3/Gag-YFP or 19RA/25KA pNL4-3 Gag-YFP. The transfected cells were incubated for 1 min at room temperature with Alexa Fluor 594-conjugated concanavalin A (ConA-AF594; Invitrogen) for visualization of the plasma membrane at 14 h post-transfection. The cells were then fixed with 4% paraformaldehyde in phosphate-buffered saline (PBS) for 30 min at room temperature, mounted with Fluoromount G (ThermoFisher) and observed with a spinning-disk confocal microscope (Nikon CSU-X1). Z-series of images were acquired with 0.2 µm intervals between the focal planes.

### Quantification and Statistical analysis

The intensity of the bands in immunoblot analysis was measured using Fiji ImageJ software (W. S. Rasband, ImageJ, U.S. National Institutes of Health, Bethesda, MD, USA, https://imagej.net/ij/, 1997–2018). Statistical analyses were performed using Prism 10 (GraphPad Software Inc., USA). The P values for statistical difference between different conditions were determined using paired Student’s t test or analysis of variance (ANOVA) and one-way Tukey’s multiple comparison test, as described in figure legends. All figures were prepared using Microsoft powerpoint and Adobe illustrator. Schematic illustrations were created with Biorender.com. Two-dimensional illustrations were made using R2DT web server (*79*).

### Accession number

The accession number of the hydro-tRNAseq and PARCLIP-hydro-tRNAseq data reported in this study is deposited to NCBI Gene Expression Omnibus (GEO) with accession number GSE314757.

## Acknowledgments

We acknowledge support from the Advanced Genomics, Bioinformatics, and Microscopy Cores of the University of Michigan Medical School’s Biomedical Research Core Facilities and support from the University of Michigan Center for Structural Biology (CSB). The CSB is grateful for support from the U-M Life Sciences Institute, the U-M Rogel Cancer Center, the U-M Medical School Endowment for Basic Sciences, and grants from the National Institute of Health. We thank Drs. Sebla Kutluay and Markus Hafner for advice on PARCLIP-hydro-tRNAseq approaches and Drs. Nicole Koropatkin, Haley Brown, and Jennifer Meagher for advice and support regarding the protein purification procedures. We also thank Dr. Karin Musier-Forsythe and members of the Ono lab for helpful discussions. Research reported in this manuscript was supported by the National Institute of Allergy and Infectious Diseases of the National Institutes of Health under Award Number R37AI071727. The content is solely the responsibility of the authors and does not necessarily represent the official views of the National Institutes of Health.

## Author contributions

Conceptualization: RM, DT, K-NC, and AO. Methodology: RM, DT, K-NC, HQ, and AO. Investigation: RM, DT, K-NC, and AO. Supervision: AO. Writing—original draft: RM and AO. Writing—review and editing: RM, DT, KC, HQ, and AO.

## Data and materials availability

All data needed to evaluate the conclusions in the paper are present in the paper and/or the Supplementary Materials.

