## Supplemental Material for "The specific interface for selenocysteine-decoding tRNA in HIV-1 Gag MA domain regulates progeny virion infectivity"

Rajat Mudgal *et al.*

**This PDF file includes:**

Fig. S1 to S8
Table S1

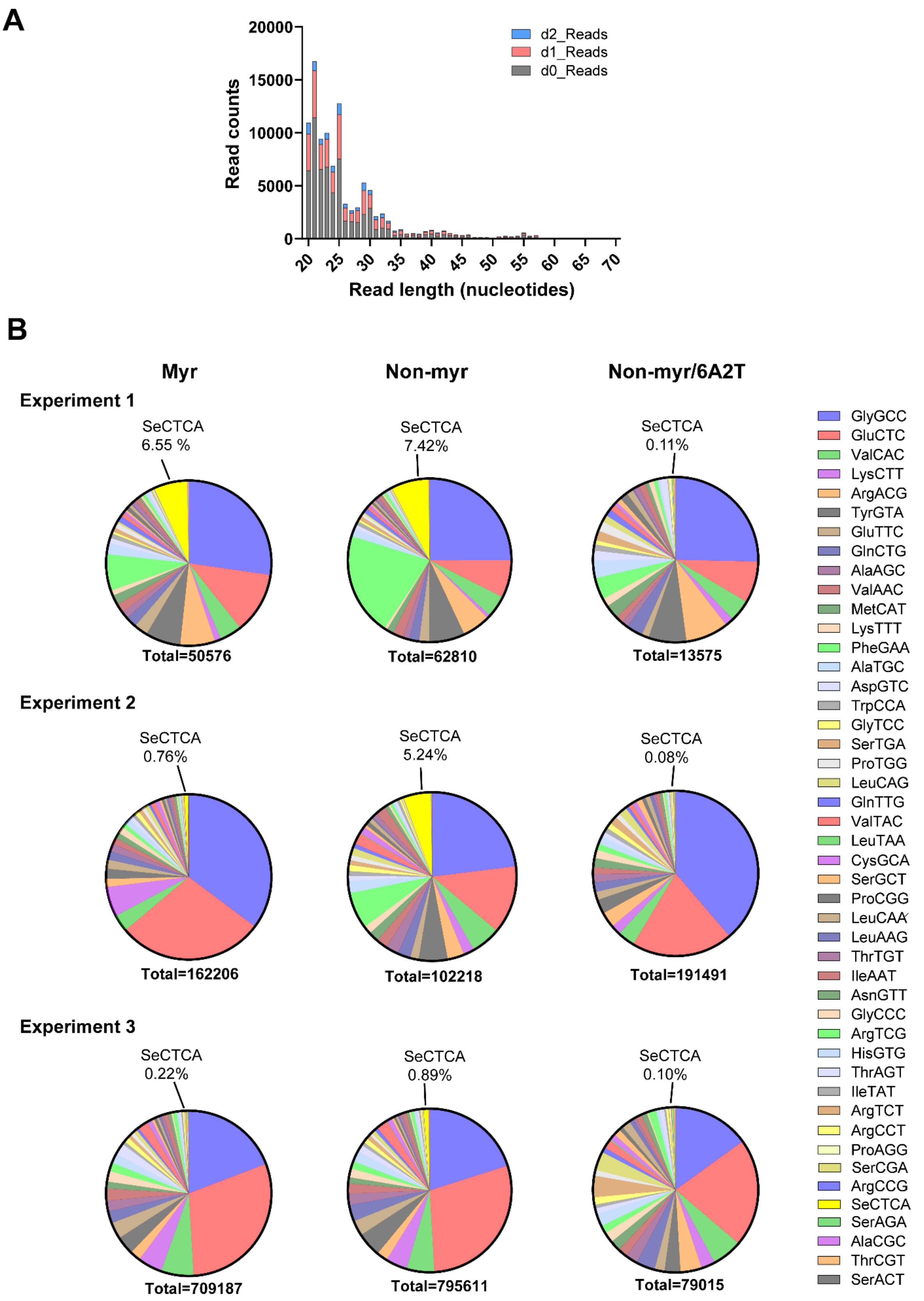


**Figure S1**

**Characteristics of reads obtained in PARCLIP-hydro-tRNAseq analysis. (A)** Representative length distribution of all adapter-trimmed reads mapped to tRNA in the reference database with 0, 1, or 2 mismatches (d0, d1, and d2). The data analyzed here are for reads of tRNA-sized RNAs bound to non-myr delNC Gag-YFP in Experiment 2. **(B)** Pie charts illustrating the percentages of different tRNAs bound to delNC Gag-YFP variants (myr, non-myr, non-myr/6A2T) in the three experiments of PARCLIP-hydro-tRNAseq shown in Figures 1 and S2. Total read counts and the percentages for SeCTCA are shown.

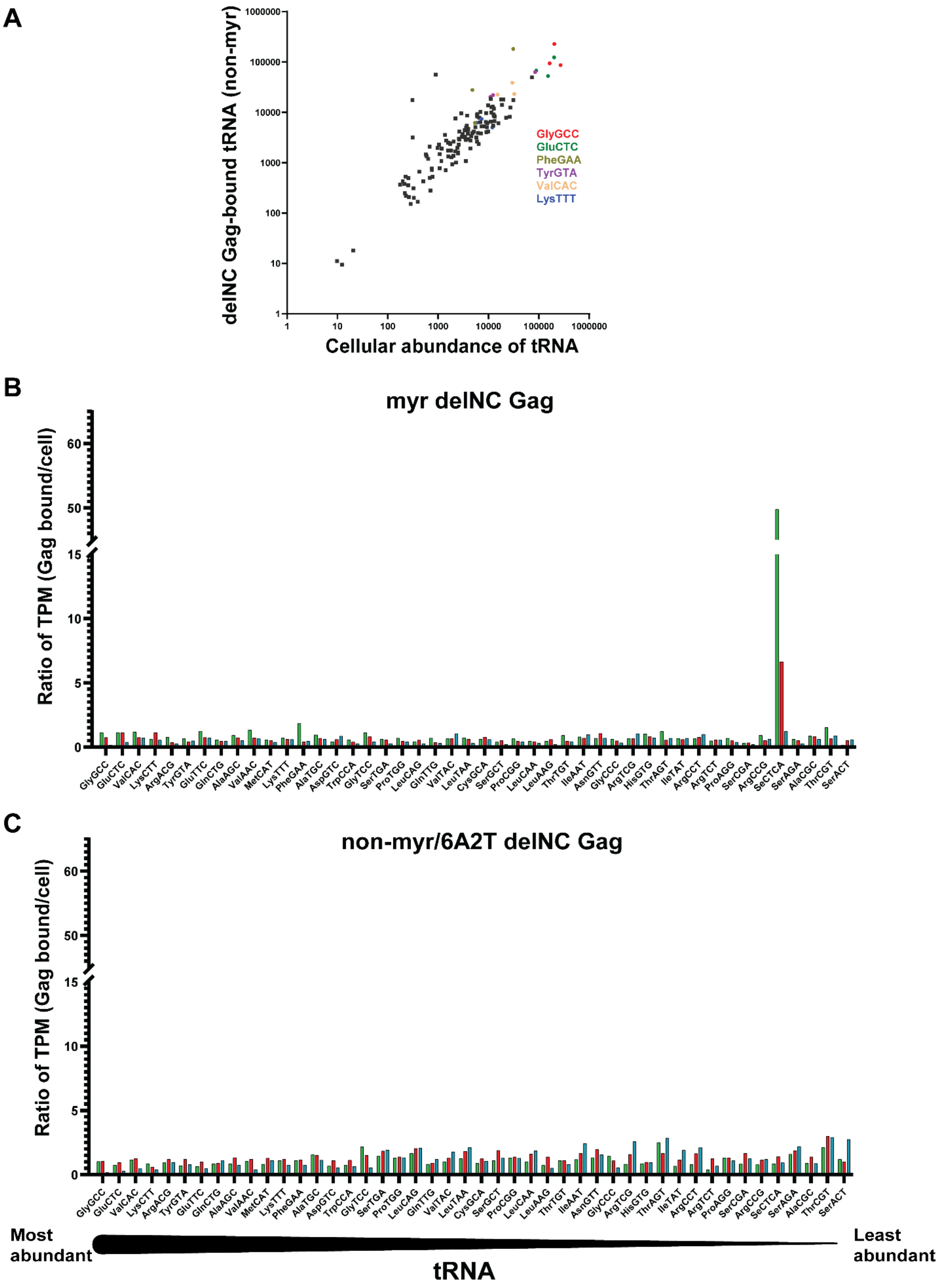


**Figure S2**

**PARCLIP-hydro-tRNAseq of Gag-bound tRNA**. **(A)** Number of reads corresponding to tRNAs present in non-myr delNC Gag-bound fraction versus tRNA abundance in HEK293T cells. Non-SeCTCA tRNAs including GlyGCC (red), LysTTT (blue), GluCTC (dark green), PheGAA (olive green), TyrGTA (magenta), and ValCAC (light orange), which are analyzed in Figures 2, 3, 4, and 6, are shown in circular dots. Data from three experiments are shown. **(B and C)** Ratios of read counts (TPM) for delNC Gag-bound tRNA and total tRNA isolated from HEK293T cells transfected with myr delNC Gag-YFP (B) or non-myr/6A2T delNC Gag-YFP (C) are plotted. tRNAs are plotted on X-axis in the decreasing order of their cellular abundance as in Figure 1C. Data from three independent experiments are shown in green, red, and blue bars. SeCTCA enrichment was observed with myr delNC Gag-bound pool of tRNAs.

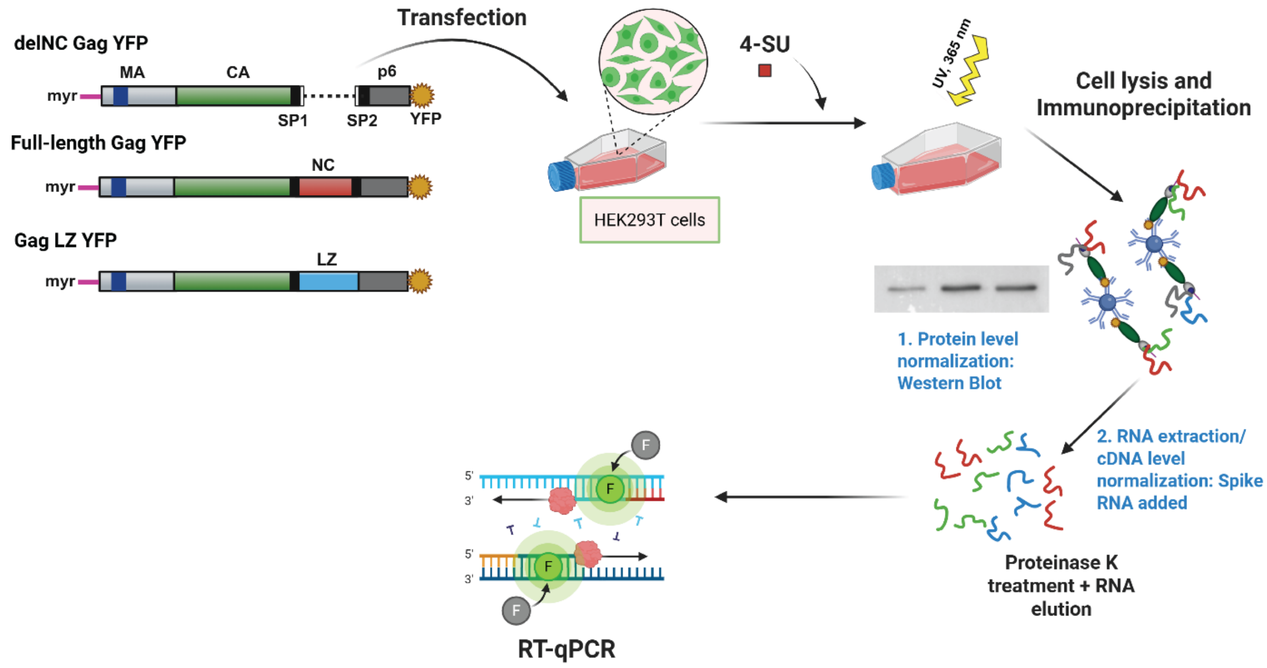


**Figure S3**

**Schematic illustration of the PARCLIP-RT-qPCR assay.** HEK293T cells were transfected with plasmids encoding delNC Gag-YFP, Full-length Gag-YFP, or GagLZ-YFP constructs containing different MA sequences (myr, non-myr, and non-myr/6A2T) or pUC19, 4-thiouridine (4-SU) was added in the culture, and RNAs containing 4-SU were crosslinked to their associated proteins with UV light. Subsequently, cell lysates were subjected to immunoprecipitation using an anti-GFP nanobody to isolate Gag. A fraction of proteins recovered after immunoprecipitation was run on SDS-PAGE and quantified after immunoblotting to measure relative levels of Gag proteins bound to nanobodies. RNA was dissociated from the recovered proteins by proteinase K treatment. An in vitro synthesized spike RNA was added to the samples prior to protease digestion to control for variations in RNA recovery and cDNA synthesis. cDNA was synthesized followed by protease digestion and subsequent phenol-chloroform extraction of RNA. RT-qPCR was performed with tRNA- and Spike RNA-specific primers. Gag protein levels and Spike RNA Cq values were used for normalizing Cq values of Gag-bound tRNA in RT-qPCR.

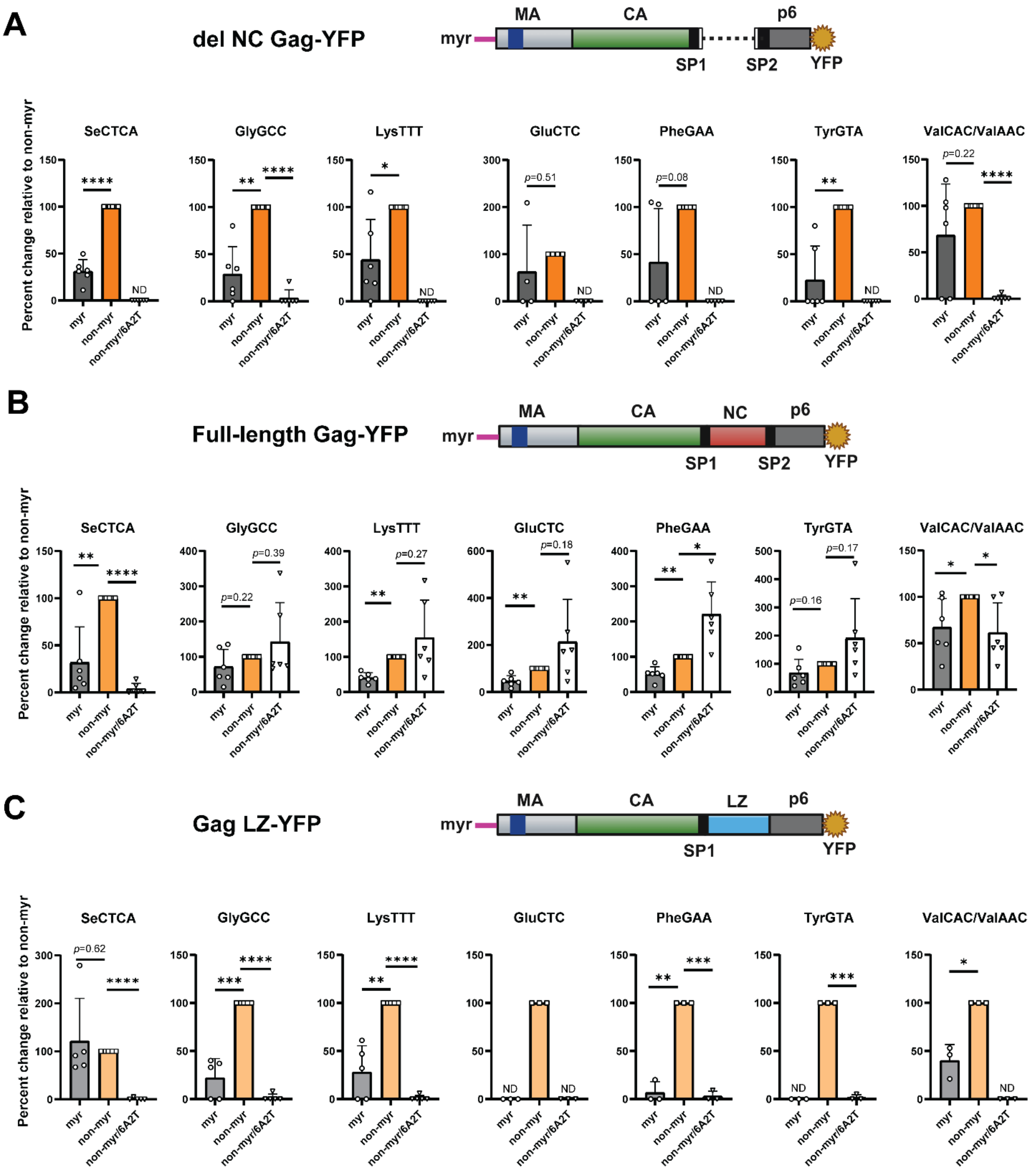


**Figure S4**

**Non-SeCTCA tRNAs bind to NC in addition to MA.** A subset of tRNAs crosslinked to Gag-YFP were quantified using PARCLIP-RT-qPCR as described in Fig. 2. tRNAs bound to cytomegalovirus (CMV) promoter-driven (A) delNC Gag-YFP (B) full-length Gag-YFP, and (C) GagLZ-YFP. LZ: Leucine Zipper motif. myr and non-myr/6A2T full-length Gag were compared with non-myr Gag for indicated tRNAs. Protein levels and Spike RNA recovery were used for normalizing Cq values in RT-qPCR. Cq values higher than or within 0.6 of the background were assigned zero**.** If data from all experiments are zero for one condition, it is indicated as ND (not detected). If the value for a given tRNA bound to non-myr Gag is zero (when binding of a tRNA to non-myr Gag did not reach the level higher than the background), we omitted the data set for that tRNA, since in this analysis we examined tRNA binding to the two other Gag constructs in comparison to the non-myr Gag. Therefore, comparisons for some tRNAs were performed with fewer (but at least three) data sets than the rest of tRNAs. Otherwise, the means ± SD of six (A-B) or five (C) independent experiments are shown. P value was determined using paired Student's t test. *P < 0.05; **P < 0.01; ***P < 0.001, ****P < 0.0001. For non-significant comparisons, p values are shown.

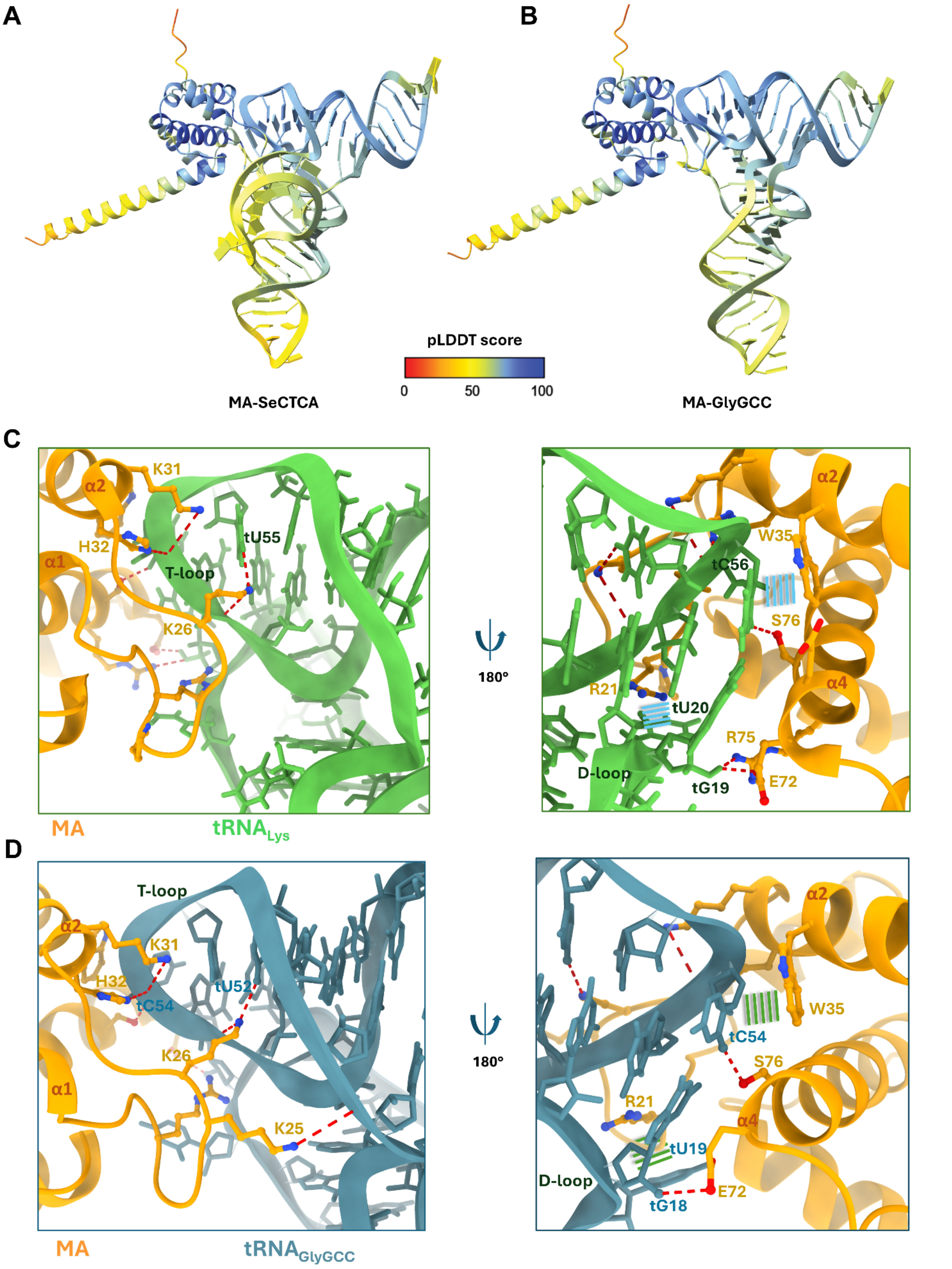


**Figure S5**

**Structural prediction and analysis of MA-tRNA interaction**. **(A and B)** The structural models of MA-SeCTCA (A) and MA-GlyGCC (B) complexes were predicted using AlphaFold 3. Local-distance difference test (pLDDT) scale is indicated, from the lowest (red, 0) to the highest confidence (blue, 100). **(C)** The interactions between MA and tRNA in the previously determined crystal structure (PDB ID: 7mrl). MA and modified $tRNA_{Lys}$ are shown in orange and green, respectively. Hydrogen bonds are shown as dashed red lines; stacking interactions as parallel blue lines. **(D)** The interactions between MA and GlyGCC in the structure of the AlphaFold 3-predicted complex. MA and GlyGCC are shown in orange and blue grey, respectively. Hydrogen bonds are shown as dashed red lines; stacking interactions as parallel green lines. Nucleotides in tRNAs are numbered with a "t" prefix.

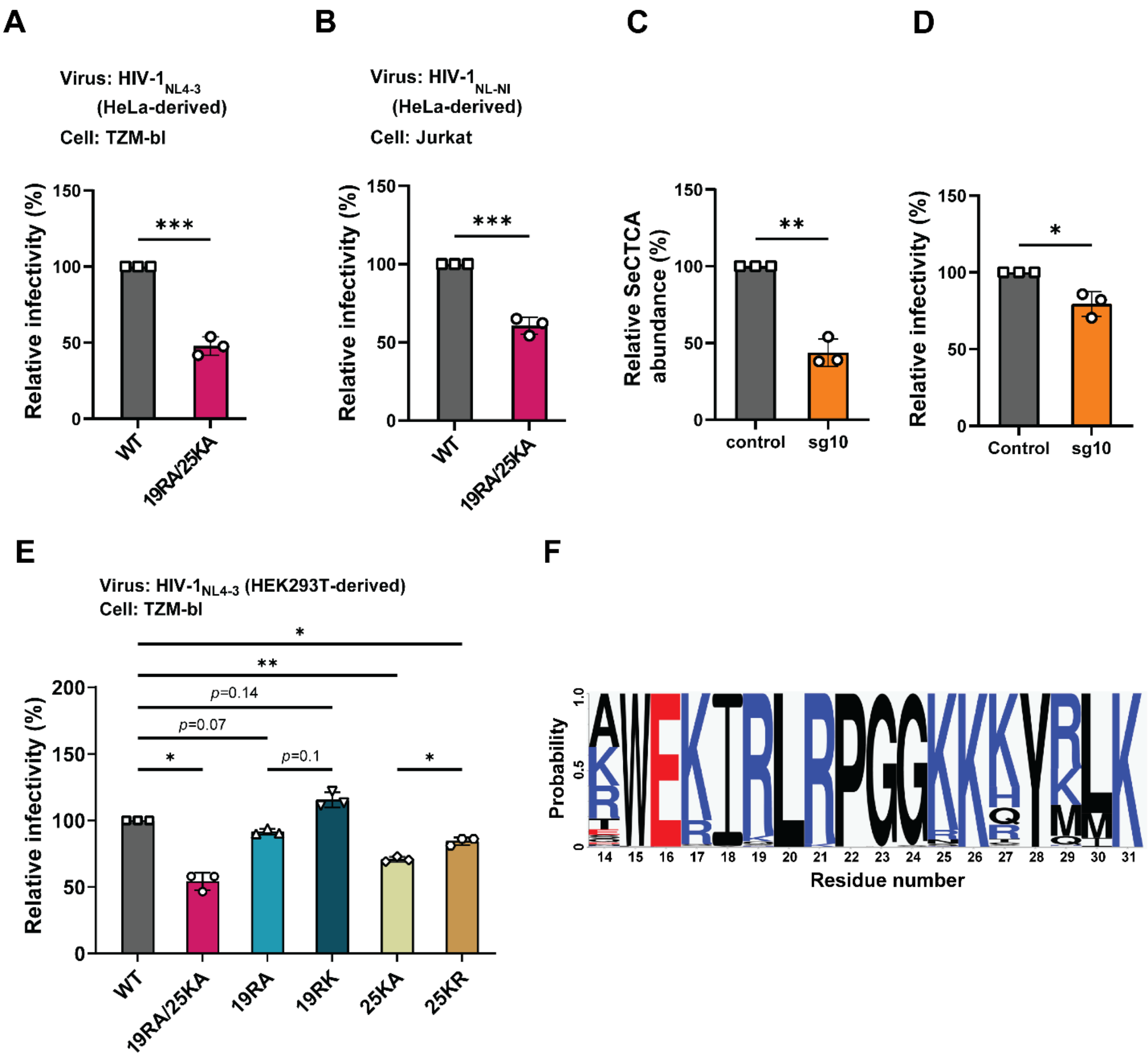


**Figure S6**

**MA-SeCTCA interaction is essential for viral infectivity. (A-B)** For infectivity analysis, HeLa cells were transfected with pNL4-3 or pNL-NI, a replication-competent HIV-1 molecular clone encoding NanoLuc, encoding WT or 19RA/25KA MA. At 16 h post-transfection, supernatants were collected and virion infectivity analysis was performed using TZM-bl cells (A) or Jurkat cells (B) as in Figure 7C-D. **(C)** Single-guide RNA-mediated knockdown (KD) of SeCTCA in HEK293T cells using CRISPR-Cas9. Relative abundance of SeCTCA in the cells was analyzed by RT-qPCR. PX459 V2.0 empty vector served as a control. **(D)** Normalized relative single-cycle infectivity of HIV-1 produced in control or SeCTCA-KD (sg10) cells was analyzed by TZM-bl assay. Luciferase activity was quantified and processed as in Figure 7C. **(E)** HEK-293T cells were transfected with pNL4-3 encoding WT MA or indicated MA variants. At 16 h post-transfection, supernatants were collected, and virion infectivity analysis was performed using TZM-bl cells as in Figure 7C. **(F)** WebLogo representation of the conservation of the MA highly basic region sequences among HIV-1 Groups M, N, O, and P as well as SIVcpz and SIVgor, generated using the filtered web sequence database at https://www.hiv.lanl.gov. All experiments were performed three times independently. Data are presented as mean ± SD. P value was determined using Student's t test or analysis of variance (ANOVA) and one-way Tukey's multiple comparison test. *$P < 0.05$, **$P < 0.01$, ***$P < 0.001$.

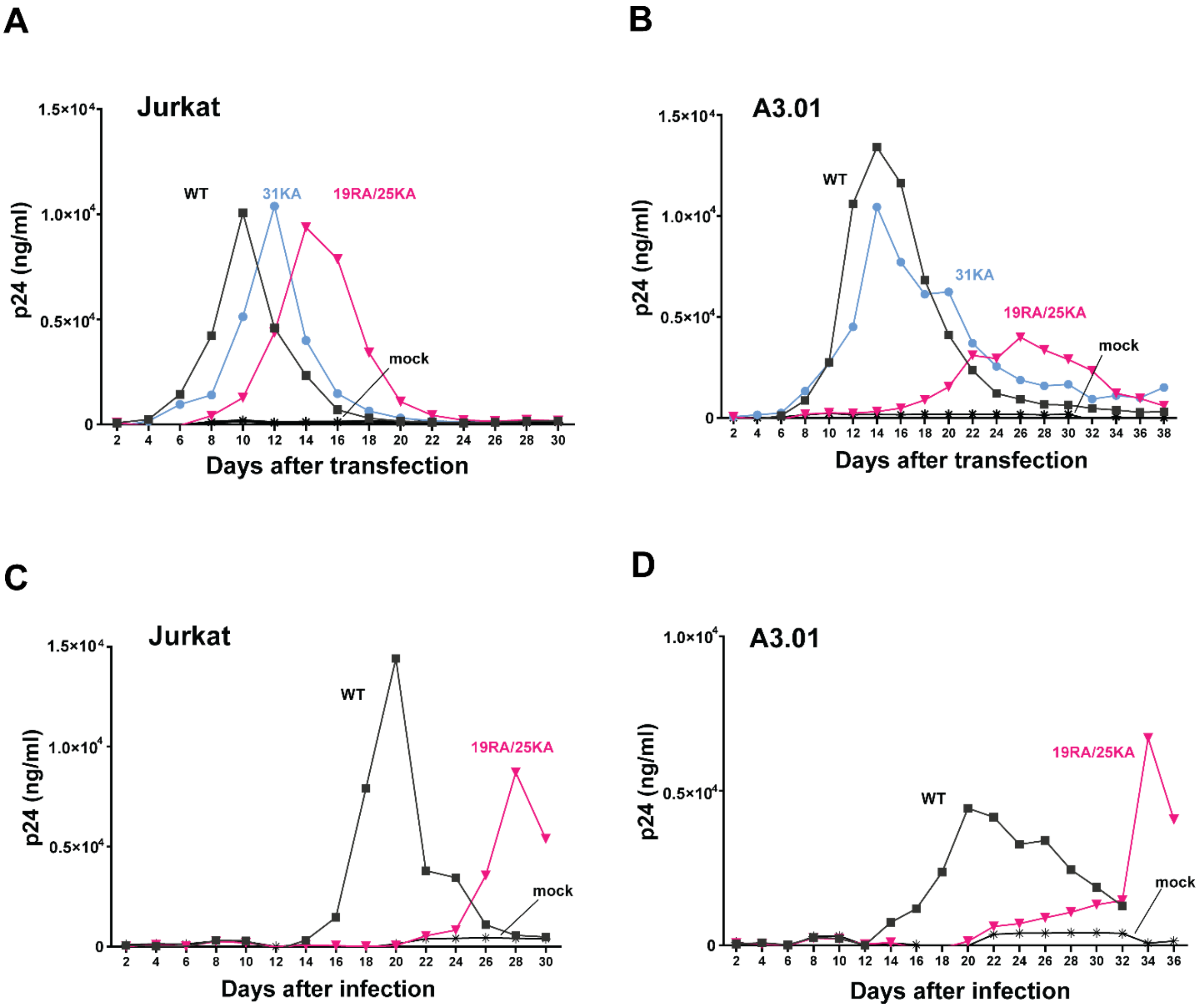


**Figure S7**

**Loss of MA-SeCTCA interaction interface impede viral replication in T cell line cultures. (A-B)** Jurkat (A) and A3.01 (B) cultures were transfected with HIV-1 molecular clones that encode WT MA, 31KA MA, or 19RA/25KA MA using DEAE-dextran. The experiments were independent repeats of those shown in Figure 8A-B. **(C-D)** Jurkat (C) and A3.01 (D) cultures were inoculated with 2.5 ng (in p24) HIV-$1_{NL4-3}$ or 5 ng 19RA/25KA HIV-$1_{NL4-3}$. Two-fold more inoculum based on p24 was used for the latter to compensate for the reduced infectivity of the inoculum. Cells were split every 2 days, and virus spread was monitored by measuring p24 released into the supernatants using p24 ELISA.

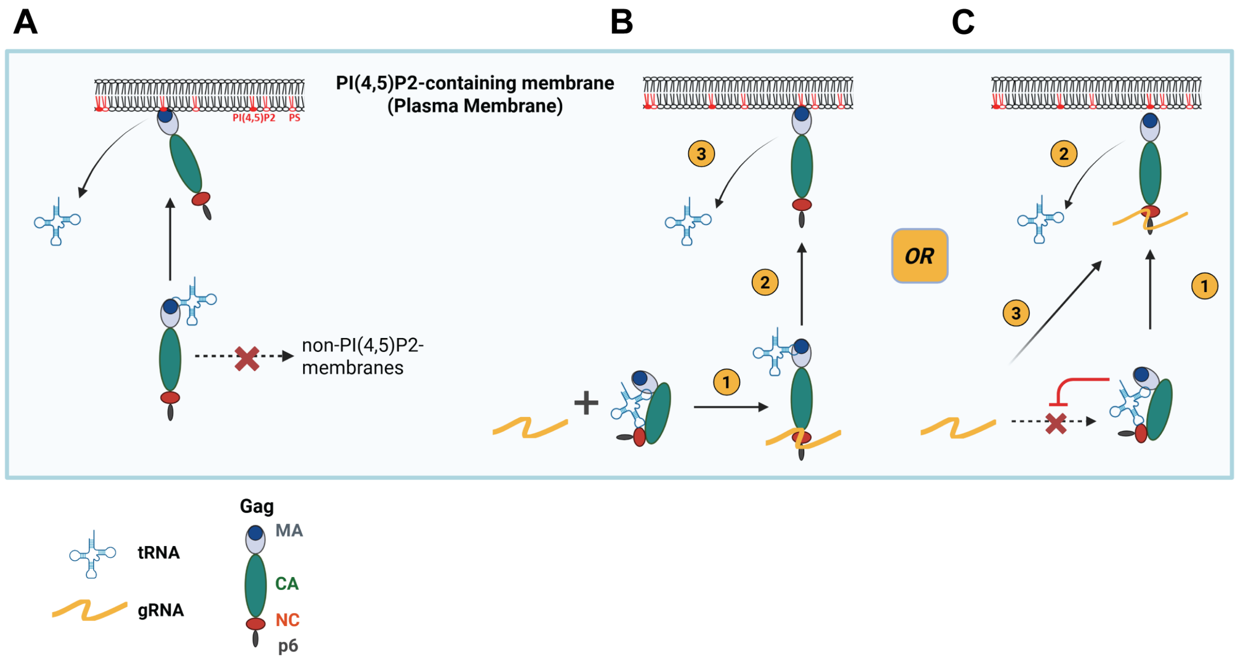


### Figure S8

**Schematic models for putative roles of MA-SeCTCA binding in HIV assembly. (A)** Binding of Gag to tRNAs blocks binding of Gag to intracellular membranes that contain phosphatidylserine (PS) but not phosphatidylinositol 4,5-bisphosphate (PIP2). Interaction of Gag with PIP2-containing membrane overcomes the block on membrane binding by tRNA, thereby recruiting Gag to the PM. **(B and C)** Hypothetical roles for tRNA binding to MA in the context of NC binding to viral RNA. (B) Non-SeCTCA tRNAs bind Gag in a bended conformation via both MA and NC domains. Binding of NC with viral genomic RNA (gRNA) in cytosol leaves the tRNA bound only to MA (1), and these Gag molecules can readily bind the plasma membrane (PM) (2) upon removal of tRNAs from MA by PIP2 (3). (C) Alternatively, non-SeCTCA tRNAs bind Gag via both MA and NC domains preventing NC from binding to genomic RNA. Upon Gag binding to the PM (1) and removal of the tRNA (2), NC becomes available for gRNA binding (3).

### Table S1.

| Table 1: Oligonucleotides used in this study | |
|---|---|
| Spike RNA | GAACCGAUUGUGGGUCAUGUAUGCCAAAGAUUGAGCCUG CUGGCAGAUGAUAGCUUCAACGAUCAUCAGUGGCCGAG |
| **Primer sets for RT-qPCR** | |
| Forward primer- GluCTC | CCCTGGTGGTCTAGTGGTTA |
| Reverse primer- GluCTC | TCCCTGACCGGGAATCG |
| Forward primer- GlyGCC | GGTGGTTCAGTGGTAGAATTCTCG |
| Reverse primer- GlyGCC | GGCCGGGAATCGAACCC |
| Forward primer- LysTTT | GTCGGTAGAGCATCAGACTT |
| Reverse primer- LysTTT | CCTGGACCCTCAGATTAAA |
| Forward primer- PheGAA | TGGGAGAGCGTTAGACTGAA |
| Reverse primer- PheGAA | AACCAGGGACCTTTAGATCT |
| Forward primer- SeCTCA | CGGATGATCCTCAGTGGTCT |
| Reverse primer- SeCTCA | GCCCGAAAGGTGGAATTGA |
| Forward primer- TyrGTA | CCTTCGATAGCTCAGCTGGTAG |
| Reverse primer-TyrGTA | GGTCCTTCGAGCCGGAAT |
| Forward primer- ValCAC/ValAAC | TAGTGTAGTGGTTATCACG |
| Reverse primer- ValCAC/ValAAC | GCCCGGTTTCGAACCGGGGAC |
| Forward primer- Spike RNA | GAACCGATTGTGGGTCATGTA |
| Reverse primer- Spike RNA | CGGCCACTGATGATCGTTGA |
| **Primers to amplify viral genome** | |
| Forward primer- 5'LTR | CCAGGGCCAGGGATCAGATATCC |
| Reverse primer- *vif* | CCTAGTGGGATGTGTACTTCTGAACTTA |
| Forward primer- *pol* | CTGGCATGGGTACCAGCACACAA |
| Reverse primer- 3'LTR | GGTCTAACCAGAGAGACCCAGTACAG |
| **Oligonucleotides to in vitro transcribe tRNAs** | |
| Forward primer- SeCTCA | GgaattcCtaatacgactcactataGCCCGGATGATCCTCAGTGGT |
| Reverse primer- SeCTCA | mUmGGCGCCCGAAAGGTGGAATTGA |
| Forward primer- GlyGCC | GgaattcCtaatacgactcactataGCATGGGTGGTTCAGTGGTAG |
| Reverse primer- GlyGCC | mUmGGTGCATGGGCCGGGAATCGAACCCGG |
| Forward primer- LysTTT | GgaattcCtaatacgactcactataGCCCGGATAGCTCAGTCGGT |
| Reverse primer- LysTTT | mUmGGCGCCCGAACAGGGACTTGAACCC |

| | |
|---|---|
| Oligo1- delV$_{12}$ SeCTCA | GgaattcCtaatacgactcactataGCCCGGATGATCCTCAGTGGTCTGGGGTGCAGGC TTCAAACCTGTAGCGAGTGGTTCAATTCCACCTTTCGGGCGCCA |
| Oligo2- delV$_{12}$ SeCTCA | mUmGGCGCCCGAAAGGTGGAATTGAACCACTCGCTACAGGTTTGAAGCC TGCACCCCAGACCACTGAGGATCATCCGGGCtatagtgagtcgtattaGgaattcC |
| Oligo1- delV$_{10}$ SeCTCA | GgaattcCtaatacgactcactataGCCCGGATGATCCTCAGTGGTCTGGGGTGCAGGC TTCAAACCTGTAGCTAGAGTGGTTCAATTCCACCTTTCGGGCGCCA |
| Oligo2- delV$_{10}$ SeCTCA | mUmGGCGCCCGAAAGGTGGAATTGAACCACTCTAGCTACAGGTTTGAAG CCTGCACCCCAGACCACTGAGGATCATCCGGGCtatagtgagtcgtattaGgaattcC |
